# Sex differences in age- and performance-related patterns of brain activity during episodic encoding and retrieval of spatial contextual details

**DOI:** 10.1101/535617

**Authors:** Sivaniya Subramaniapillai, Sricharana Rajagopal, Abdelhalim Elshiekh, Stamatoula Pasvanis, Elizabeth Ankudowich, M. Natasha Rajah

## Abstract

Aging is associated with episodic memory decline and alterations in memory-related brain function. However, it remains unclear if age-related memory decline is associated with similar patterns of brain aging in women and men. In the current task fMRI study, we tested the hypothesis that there are sex differences in the effect of age and memory performance on brain activity during episodic encoding and retrieval of face-location associations (spatial context memory). Forty-one women and 41 men between the ages of 21 to 76 years participated in this study. Between-group multivariate partial least squares (PLS) analysis of the fMRI data was conducted to directly test for sex-group differences and similarities in age-related and performance-related patterns of brain activity. Our behavioural analysis indicated no significant sex differences in retrieval accuracy on the fMRI tasks. In relation to performance effects, we observed similarities and differences in how retrieval accuracy related to brain activity in women and men. Both sexes activated dorsal and lateral prefrontal cortex (PFC), inferior parietal cortex (IPC) and left parahippocampal gyrus (PHG) at encoding and this supported subsequent memory performance. However, there were sex differences in retrieval activity in these same regions and in lateral occipital-temporal and ventrolateral PFC. In relation to age effects, we observed sex differences in the effect of age on memory-related activity within PFC, IPC, PHG and lateral occipital-temporal cortices. Overall, our findings suggest that the neural correlates of age-related spatial context memory decline differ in women compared to men.

## 1. Introduction

Episodic memory is our ability to encode, store and retrieve personally experienced events in rich contextual detail (Tulving, 1984). The ability to encode and retrieve contextual details of episodic memories diminishes with age and impacts adult quality of life (e.g., Cansino, 2009; Dulas & Duarte, 2012; Glisky, Polster, & Routhieaux, 1995; Rajah, Languay, & Valiquette, 2010; Schacter, Kaszniak, Kihlstrom, & Valdiserri, 1991; Spencer & Raz, 1995; Trott, Friedman, Ritter, & Fabiani, 1997). Neuroimaging studies have shown that encoding and recollecting item-context associations relies on the function of a distributed network of brain regions including the medial temporal lobe (MTL), prefrontal cortex (PFC), and inferior parietal cortex (IPC) (Grady et al., 2010; Hayama, Vilberg, & Rugg, 2012; Johnson & Rugg, 2007; Rajah, Crane, Maillet, & Floden, 2011; Rajah, McIntosh, & Grady, 1999; Spaniol et al., 2009). As such, age-related declines in episodic memory for contextual details (context memory) have been attributed to structural and functional differences in these brain regions. For example, in our previous adult lifespan fMRI study, we found that age-related deficits in context memory started in early midlife and was associated with differences in bilateral fusiform gyrus, anterior PFC and ventrolateral PFC activity in middle-aged, compared to young, adults (Kwon et al., 2016). In older adults context memory decline was more severe and was associated with differences in dorsolateral PFC, MTL and IPC activity in older adults, compared to middle-aged and young adults (Ankudowich, Pasvanis, & Rajah, 2016, 2017). Previous fMRI studies have also reported age-related differences in context memory with corresponding functional differences in PFC, MTL, and parietal areas (Cansino, Estrada-Manilla, et al., 2015; Cansino, Trejo-Morales, et al., 2015; Cansino, Hernández-Ramos, & Trejo-Morales, 2012; Cansino, Trejo-Morales, & Hernández-Ramos, 2010; Dennis et al., 2008; K. J. Mitchell, Ankudowich, Durbin, Greene, & Johnson, 2013, but see Cabeza, Anderson, Locantore, & McIntosh, 2002 for functional similarities in context memory with aging). Therefore, there is growing consensus that age-related declines in context memory for visually presented stimuli are linked to differences in frontoparietal, medial temporal and ventral visual function with age.

However, the majority of fMRI studies that have investigated age-related differences in brain activity during episodic memory have pooled male and female participants. This assumes the neural correlates of age-related episodic memory decline are similar in women and men. Yet, there is evidence for sex differences in brain structure and function in adulthood (see Gur & Gur, 2002 for review), and evidence that women and men may perform differently on specific episodic memory tasks, depending on the experimental stimuli and design. For example, behavioural studies demonstrate that females perform better on episodic memory tasks for verbal stimuli (Gur & Gur, 2002; Herlitz, Nilsson, & Bäckman, 1997; Kimura & Harshman, 1984; Ragland, Coleman, Gur, Glahn, & Gur, 2000), negative emotional stimuli (Young, Bellgowan, Bodurka, & Drevets, 2013), face stimuli (Keightley, Winocur, Burianova, Hongwanishkul, & Grady, 2006; Sommer, Hildebrandt, Kunina-Habenicht, Schacht, & Wilhelm, 2013; J. E. Yonker, Eriksson, Nilsson, & Herlitz, 2003), and verbal paired associated memory (Bender, Naveh-Benjamin, & Raz, 2010), compared to men. In contrast, on average, males have performed better than females on object-location associative memory tasks (Postma, Izendoorn, & De Haan, 1998) and memory tasks assessing visuospatial ability (e.g., De Frias, Nilsson, & Herlitz, 2006; Weiss, Kemmler, Deisenhammer, Fleischhacker, & Delazer, 2003).

Exploring the origins of sex differences in brain aging and episodic memory function is important clinically, since studies have reported sex differences in the incidence, and risk, for memory-related disorders (Azad, Al Bugami, & Loy-English, 2007; Jorm & Dolley, 1998). For example, women are more at risk of developing late-onset Alzheimer’s Disease, compared to men (Alzheimer’s Association, 2015; Mielke, Vemuri, & Rocca, 2014). Therefore, it is critical to develop a working model of healthy aging in both sexes in order to understand how the interaction of biological sex and age can lead to pathological aging i.e., AD.

However, a feature of, sex differences in episodic memory and brain structure/function is that sociological and environmental factors influence brain development and cognitive/behavioural strategic preferences between the sexes. Thus, sex differences in brain structure and function may be observed, *even when sex differences in cognitive/behavioural performance are not*. For example, Nyberg et al (2000) found both similarities and differences between sexes when they investigated sex differences in the neural correlates of episodic memory retrieval in a younger adult sample (17 men; 17 women) (Nyberg, Habib, & Herlitz, 2000). In this study, subjects responded yes/no in a recognition paradigm for words, sentences, and landscape images. No sex differences in recognition memory were reported. A multivariate partial least squares (PLS) analysis of the O^15^ positron emission tomography (PET) data revealed both similarities and differences in memory-related brain activity in women and men. Both women and men exhibited retrieval-related activity in right PFC, anterior cingulate gyrus, and midline occipital-parietal areas, which are regions typically recruited during episodic retrieval (Cabeza & Nyberg, 2000; Lepage, Ghaffar, Nyberg, & Tulving, 2000). In addition, sex differences in retrieval-related activity were observed: women showed greater retrieval-related activity in the anterior cingulate gyrus, right inferior frontal gyrus, right fusiform, parietal cortex and cerebellum compared to men. Men showed greater activity compared to women at retrieval in bilateral inferior frontal cortex (BA44, 45), bilateral inferior temporal and parietal cortex, and in posterior cingulate cortex. Similarly, previous work have reported functional sex differences during memory tasks, albeit with a greater focus on investigating autobiographical memory (St Jacques, Conway, & Cabeza, 2011; Young et al., 2013)

Importantly, sex *and* age differences in PFC, parietal, MTL and/or ventral occipitotemporal function during episodic memory studies have also been reported (Compère et al., 2016; Haut & Barch, 2006; Ragland et al., 2000; Spalek et al., 2015). Yet, it remains unclear if biological sex influences the impact of age on the neural correlates of episodic memory. In other words, we do not know if there is a sex-by-age interaction, and whether the results from prior neuroimaging studies of aging and episodic memory equally reflect how age impacts brain function and episodic memory in women and men, or if some results were driven by one sex. Moreover, most prior studies have been limited in terms of specifically testing autobiographical or recognition memory, and in terms of the age range of participants used (i.e., only young and/or middle-aged participants) (Compère et al., 2016; Haut & Barch, 2006; Ragland et al., 2000; Spalek et al., 2015). Given that sex differences in episodic memory remain relatively stable across the adult lifespan (De Frias et al., 2006; Jack et al., 2015), one may hypothesize that sex differences in the functional neural correlates of episodic memory, if present, are stable across the adult lifespan into older age. However, there is a growing body of human neuroimaging studies showing sex hormone levels in women and men differ across the lifespan, and accordingly affects brain function and cognition. For example, menopause-related decline in 17-beta-estradiol (e.g., Jacobs et al., 2016b; Morrison, Brinton, Schmidt, & Gore, 2006; Rentz et al., 2017) or age-related decline in testosterone (e.g., Moffat et al., 2018; Muller, Aleman, Grobbee, de Haan, & van der Schouw, 2005; Rosario, Chang, Head, Stanczyk, & Pike, 2011; Rosario & Pike, 2008), may contribute to differences in cognitive function. In addition, sociocultural factors, such as self-perceived gender trait possessions (Hamilton, 1995; Sharps, Price, & Williams, 1994), may also influence task performance (Baenninger & Newcombe, 1989; Hamilton, 1995; Lawton, 1994; Sharps et al., 1994). Therefore, it is also possible that there are sex differences in how age impacts brain function and episodic memory. This implies that the functional neural basis of age-related memory decline may differ in women, compared to men. Such an observation would have important clinical implications since it would suggest that specific interventions may be required in women, compared to men, to support the maintenance of memory function in late life. We test this possibility in the current study.

The goal of the current cross-sectional fMRI study is to investigate how biological sex impacts the neural correlates of episodic memory encoding and retrieval of face-location association (spatial context memory) in an adult lifespan sample. A subsample of age- and education-matched women and men from a previously collected fMRI data sample were used (Ankudowich et al., 2016). Due to prior studies highlighting the impact of menopausal transition on episodic memory-related brain regions PFC and MTL (Jacobs et al., 2016a, 2016b; Rentz et al., 2017), and our limited sample size for this subsample (N = 10), we excluded middle-aged women who self-reported being in menopausal transition or being in the early stages of postmenopause in the current study. We used a between group multivariate behaviour partial least squares (B-PLS) statistical analysis to examine sex differences in the effect of age and memory performance on brain activity during episodic encoding and retrieval (McIntosh et al., 2004). The rationale for choosing this method of analysis, in comparison to more traditional univariate contrast-based approaches, was that we wanted to use a data-driven method to objectively assess if there were patterns of whole-brain activity that differentiated how age impacted memory-related brain activity in women, compared to men. Although a contrast-based method could also be used to test this hypothesis, by not using a priori contrasts we are able to observe what the strongest effects in our dataset are, and whether sex differences were one of them. Furthermore, B-PLS allows for a direct assessment of sex differences in how age, memory performance, and Age*Memory performance impacts brain activity using. Thus, it is a powerful and parsimonious multivariate method that allows researchers to capture complex relationship between whole-brain patterns of brain activity and multiple exogenous variables of interest in one mathematical step (Andersen et al., 2012; McIntosh et al., 1996). In addition, statistical significance of B-PLS experimental effects are based on permutation testing at the image level; and, the stability of brain activation patterns identified by PLS are assessed with bootstrapping, which protects against the impact of outliers (McIntosh et al., 2004). Thus, in using B-PLS for the present study, we aimed to identify functional activation patterns from fMRI data at the whole-brain level that covaried with memory performance, and to further identify how these patterns differ in women compared to men across the aging lifespan. Based on prior behavioural and fMRI studies in young adults (e.g., Nyberg, Habib, & Herlitz, 2000; Spaniol et al., 2009), we predicted there would be sex differences in brain activity during encoding and retrieval. Additionally, given the literature on age-related differences in sex hormone levels and their influence on cognition, we also predict that there will be sex differences in how age influences memory-related brain activity at both encoding and retrieval.

## 2. Materials and Methods

### 2.1. Participants

Participants were 41 men and 41 women between the ages of 21-76 years (M = 47.1, SE = 2.03). Thirty participants could be classed as younger adults (age range: 21-35, M = 26.3, SE = 0.60), 20 as middle-aged (age range: 40 – 58, 45.8, SE = 1.17), and 32 as older adults (age range: 60-76, M = 67.3, SE = 0.66). In each age group, we had an equal number of men and women that were closely matched according to age and level of education. This sample was obtained from a larger cohort sample (N = 128) primarily used for the investigation of episodic memory across the aging lifespan (e.g., Ankudowich et al., 2016). We included middle-aged women who self-reported as pre-menopausal (excluding N = 18 currently menopausal MA women), and excluded female participants who had undergone hormone replacement therapy (HRT) (excluding N = 10 females) since menopausal transition and HRT influence brain function and episodic memory abilities (Henderson, 2010; Li, Cui, & Shen, 2014; Rentz et al., 2017; Yonker et al., 2006). Finally, we matched the male sample (N = 43) to the existing subsample of females according to age and education level, in order to use a balanced sample of males and females. Analysis of the data revealed two outliers which were subsequently removed to obtain the final matched sample.

All participants had at least a high school education (mean education = 16.1 yrs, SE = 0.22). Everyone was right-handed as assessed using the Edinburgh Inventory for Handedness (Oldfield, 1971), and met the inclusion/exclusion criteria (listed below). The study was approved by the Institutional Review Board of the Faculty of Medicine, McGill University, and all participants signed a consent form prior to participating in the study. Table 1 summarizes participant demographic and neuropsychological information by Sex within each Age Group. Post-hoc analyses revealed no significant differences in sex and education level between men and women among each age group (p > 0.05). Participants were recruited via newspaper and online advertisements in Montréal, Canada.

**Table 1.**
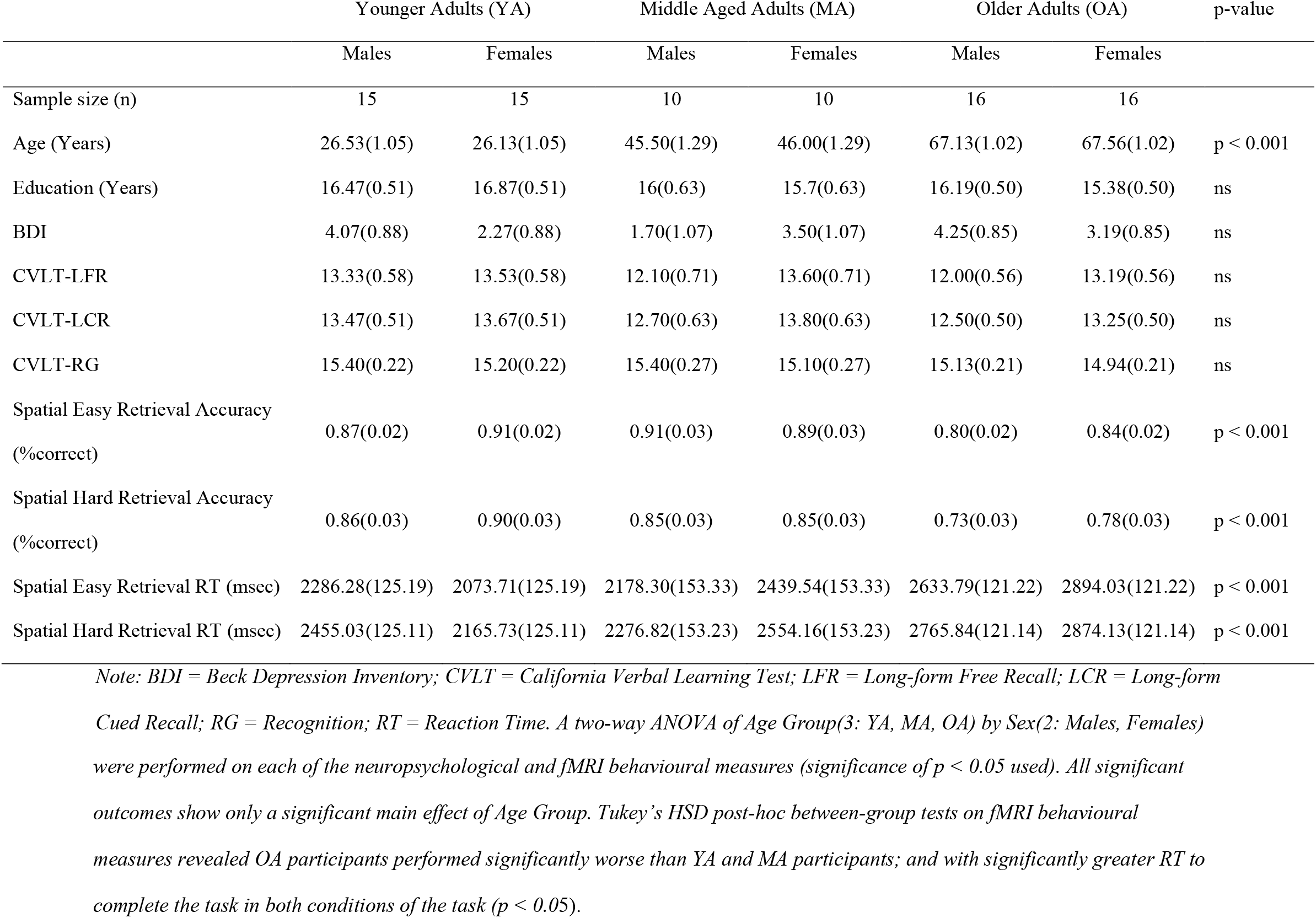
Mean Demographic and fMRI Behavioural Measures (and Standard Errors) by Age Group & Sex

### 2.2. Behavioural Methods

Experimental details of this study have been previously published (Ankudowich et al., 2016). All testing was done at the Douglas Institute Brain Imaging Center. There were two testing sessions in the experiment. The first session involved medical and neuropsychological testing to ensure participants met inclusion and exclusion criteria to participate in the study. This included completing the Mini Mental Status Exam (MMSE, exclusion cut-off score < 27, (Folstein, Folstein, & McHugh, 1975), the Beck Depression Inventory (BDI) (inclusion cut-off < 15 (Beck & Steer, 1987), the American National Adult Reading Test (NART) (inclusion cut-off ≤ 2.5 SD (Strauss, Sherman, & Spreen, 2006). Participants were additionally excluded from the study if they self-reported as having a history of neurological insult, psychiatric illness, substance abuse, smoking > 40 cigarettes/day, diabetes, BMI > 30, or were currently diagnosed with high cholesterol levels and/or high blood pressure that was untreated or had only been treated for less than six months. Participants also completed the California Verbal Learning Task (CVLT) which was used to assess verbal item memory. All the aforementioned participants met the cut-off criteria for the neuropsychological tests and were able to perform above chance on a mock fMRI session of the context memory tasks employed in our fMRI study (detailed below), and were invited to the second fMRI scanning session. The stimuli used in the mock scanning session did not overlap with stimuli used in the fMRI session.

During the fMRI Session, blood-oxygen-level-dependent (BOLD) fMRI scans were obtained while participants performed easy and difficult versions of spatial and temporal context memory tasks for photographs of age-variant face stimuli of multiple ethnicities. Scans were obtained during both encoding and retrieval phases of the memory tasks. The difficulty manipulation was included to help differentiate brain activity related to main effects of sex and performance, and Sex × Performance interactions. Details of the difficulty manipulation are presented below. A mixed rapid event-related fMRI design with 12 experimental runs was employed (Dale & Buckner, 1997). Each run contained a spatial easy (SE), temporal easy (TE), and a hard version of either a spatial (SH) or temporal (TH) memory task. In total, the SE and TE tasks were performed 12 times and the SH and TH tasks were performed six times. The results herein only examined the behavioural and fMRI data collected during spatial context memory tasks to reduce the complexity of the fMRI analysis conducted. Moreover, we focused specifically on the spatial context memory tasks to facilitate the interpretations of our findings in light of the considerable literature investigating sex differences in spatial episodic memory (Herlitz et al., 1997; 1999; 2008, Lewin et al., 2001). Thus, only details pertaining to these tasks are presented below. For details about the temporal context memory tasks, refer to Ankudowich et al. (2016; 2017). The behavioural tasks were designed and implemented using E-Prime (Psychology Software Tools, Inc.; Pittsburgh, PA, USA), and accuracy (% correct) and reaction time (RT, msec) was collected. Below we explain the experimental details for spatial context encoding and retrieval.

### 2.3. Spatial Context Encoding

Prior to each encoding session, participants were presented with a 9 sec instruction screen which informed them that the following task was a spatial context memory task, and that they were required to memorize each face and its left/right spatial location on the screen. In addition, subjects were asked to rate each face as being pleasant or neutral using a button press. We have found that making such a social-emotional evaluation of stimuli at encoding improves subsequent memory (Maillet & Rajah, 2013, but also see Harvey, Fossati, & Lepage, 2007; Mitchell, Macrae, & Banaji, 2004). After the encoding instructions were presented, participants were shown a series of faces, one at a time, for 2 sec/stimulus with a variable inter-trial interval (ITI) of 2.2-8.8 sec, which served to add jitter to the event-related fMRI data acquisition (Dale & Buckner, 1997). Each face was presented either on the left or right side of a central fixation. During SE tasks, six faces were presented at encoding and during SH tasks, 12 faces were presented. Therefore, encoding load manipulation was used to modulate task difficulty. In total, there were 72 encoding stimuli per task (12 blocks of SE encoding tasks and 6 blocks of SH encoding tasks).

Encoding was followed by a one-minute distraction phase where participants performed a verbal alphabetization task which required participants to identify which one of two words came first in the alphabet. The goal of this part of the experiment was to prevent participants from actively rehearsing the face stimuli.

### 2.4. Spatial Context Retrieval

After the distraction phase, participants entered the retrieval phase of the task. There was a 9 sec instruction screen that appeared prior to retrieval events to orient participants to their task. Participants were instructed they would see a series of two previously encoded faces during each retrieval event and were required to select which of the two faces, were originally presented on the left of the screen (for half the retrieval tasks) or on the right of screen (for half the retrieval tasks). After the instructions, participants were presented with three consecutive retrieval events, 6 sec/event, with a variable ITI (as stated above) during SE tasks, and with six consecutive retrieval events for SH tasks. Participants made their responses using a button press with an MRI-compatible response box. There was a total of 36 retrieval events per task type across the experiment.

### 2.5. MRI methods

The fMRI scanning took place at the Douglas Institute Brain Imaging Center, using a 3T Siemens Magnetom Trio scanner. Participants laid in a supine position in the scanner wearing a standard head coil. First, the T1-weighted anatomical image using a 3D gradient echo MPRAGE sequence (TR= 2300 msec, TE= 2.98 msec, flip angle = 9°, 176 1 mm sagittal slices, 1 × 1 × 1 mm voxels, FOV = 256 mm^2^, GRAPPA of 2, acquisition time = 5 min) was acquired. After the structural scan, functional BOLD MRI images were collected using a single-shot T2*-weighted gradient echo-planar imaging (EPI) pulse sequence (matrix size = 64 × 64, 32 oblique slices with no slice gap, in-plane resolution of 4×4 mm, TR = 2000 ms, TE = 30 ms, 4 mm^3^ isotropic, FOV = 256 mm^2^) while subjects performed the aforementioned behavioural task. The memory task was initiated on a computer using E-prime software, which was back-projected onto a screen in the scanner bore, and made visible to participants via a mirror mounted within the head coil. Participants were able to make task-related responses using a fiber-optic 4-button response box. Participants requiring correction for visual acuity wore corrective plastic lenses. Twelve fMRI runs were conducted, as outlined in the behavioural methods; each run was approximately 9 minutes long and yielded 278 whole-brain volumes per run. Total time in the scanner, with setup, was approximately two hours.

### 2.6. Preprocessing

Raw DICOM files were converted to ANALYZE format. The first five functional images were removed to ensure that tissue had reached a steady state magnetization. The fMRI data were preprocessed using Statistical Parametric Mapping (version 8). ArtRepair from SPM8 Toolbox was used for noise filtering to correct for bad slices and voxel spike noise using linear interpolation for up to 5% of the fMRI data and, and elimination of data outside the head. After correction for bad slices, images were realigned to the first acquired functional image and corrected for movement using a 6-parameter rigid-body spatial transform. Participants with greater than 4mm of movement within a run were removed from the analysis. None of the participants in the current study moved > 4 mm. The data were then normalized to the MNI EPI template (4×4×4 mm voxel resolution) using a 12-parameter affine transformation with default settings, and then smoothed using an 8 mm FWHM isotropic Gaussian kernel. Finally, ArtRepair (SPM8) was again used to correct for bad volumes using a standard of <5% interpolated data.

### 2.7. Behavioural Data Analysis

#### 2.7.1. Spatial Context Retrieval Accuracy & RT

Using R (R Core Team, 2016), we conducted a linear mixed effects regression (LMER) model (using the ‘lme4’ package in R; Bates et al., 2015) to test the three-way interaction between Age, Sex(2: males, females), and Task Difficulty (2: easy, hard) on retrieval accuracy (% correct) and reaction time (msec). In addition, the model contained the random effect of subject to account for the variability of subject performance between the two difficulty conditions of the spatial task. Thus, in terms of R syntax, the specific model that was fitted was:

> Spatial Accuracy ~ Age *Sex*Task Difficulty + (1 | Subject)

The variables of Age and Education were standardized using a Z-score transformation and treated as continuous variables. Sex and Task Difficulty were treated as categorical variables through deviation coding (−1, 1). Significance was computed via the Satterthwaite approximations (at p ≤ 0.05; using the R ‘lmerTest’ package, Kuznetsova et al., 2017).

### 2.8. Multivariate Behavioural-Partial Least Squares Analysis

We used a between group (i.e., Sex) multivariate Behavioural-Partial Least Squares (B-PLS; (McIntosh & Lobaugh, 2004) to assesses the relationship between event-related brain activity (as measured by fMRI), age and retrieval accuracy in females, compared to males. B-PLS is a powerful method that permits analysis of large datasets (e.g., neuroimaging data) in order to assess the spatiotemporal distribution of whole-brain patterns of task-related activity and behaviour. Importantly, this analysis does not make any assumptions about the shape of the hemodynamic response function, and makes use of permutation testing and bootstrap resampling methods to identify statistically robust multivoxel patterns of brain activity over time (McIntosh & Lobaugh, 2004).

In order to run a B-PLS analysis, first, the event-related brain activity and the behavioural measures must be stored in separate matrices. The *brain activity matrix* consists of the BOLD signal at each voxel (columns of the matrix) for each participant (rows of the matrix, ordered by event type) across the voxel time series (time lags 0-7 with a TR = 2 sec for a total of 16 seconds). Importantly, we only analyzed fMRI activity for correctly remembered events during encoding and retrieval. There were four event types, which were encoding spatial easy (eSE), encoding spatial hard (eSH), retrieval spatial easy (rSE), and retrieval spatial hard (rSH). The whole brain BOLD fMRI activity for these event-types were stacked according to sex, where male and female groups were stacked above one another. In order to create the *behavioural matrix*, the behavioural measures of age and retrieval accuracy were used. The age variable was first submitted to a regression analysis in which retrieval accuracy was used to predict age. The age residual was used because age and accuracy variables are highly correlated and using the age residual (which is uncorrelated to retrieval accuracy) would minimize collinearity of the two behavioural measures. The age residual and retrieval accuracy measures were stored in the final behavioural matrix, similarly to how the brain matrix was structured.

Both brain and behavioural matrices were cross-correlated to create the covariance matrix. This matrix was submitted to singular value decomposition (SVD), which generates latent variables (i.e., orthogonal singular vectors) that maximally captures the relationship between brain activity and behaviour. For each latent variable (LV), PLS outputs 1) a singular value, 2) a correlation profile, and 3) a singular image. The singular value represents the covariance accounted for by that particular LV; all LVs are ranked (i.e., highest to lowest) according to the amount of covariance that the singular image explains out of the brain-behaviour relation. The correlation profile shows brain-behaviour relations between brain activity and the two behavioural measures of interest (i.e., age residual and accuracy). The correlation profile shows how subjects’ age and retrieval accuracy correlates with the pattern of brain activity shown in the singular image. Finally, the singular image of brain saliences contains positive and/or negative voxel salience regions which reflects the different relationships between brain activity and the behavioural measures of interest (age, accuracy). The brain-behaviour correlation profile and brain saliences reflect a symmetrical (relationship) pairing, where the positive/negative brain saliences represent whether activity in a voxel is positively or negatively associated with the correlation profile associated with that singular image. For instance, a correlation profile with an Accuracy measure that has a positive correlation at encoding would mean that this behavioural measure is negatively correlated to brain activity in the negative voxel salience regions (blue coloured regions shown in the singular image) and positively correlated to positive voxel salience regions (warm coloured regions shown in the singular image) identified by this LV. However, if a behavioural measure (e.g., Age residual) has a negative correlation at retrieval, this would mean that this Age residual is positively correlated to brain activity in the negative voxel salience regions and negatively correlated to positive voxel salience regions identified by this LV.

Statistical significance of the LV was assessed using permutation tests (assessing LV effect) and bootstrap tests (assessing stability of voxel saliences). The significance of the LV effect was assessed using permutation testing, by testing whether a signal was significantly different from noise. Each permutation involves reassigning the fMRI data (event/condition) and behavioural measures (age, retrieval accuracy) within subject by resampling, without replacement. A PLS was rerun for each resampled dataset and the probability that the permuted singular values exceed the observed singular value for a given LV was used to determine significance at p < 0.05 (McIntosh & Lobaugh, 2004).

After permutation testing, bootstrap tests were conducted to calculate the reliability of the voxel saliences in contributing to the singular image for each LV. Thus, the bootstrap procedure identifies voxel saliences with the most reliable patterns of positive and negative brain saliences shown in the singular image. We used 500 bootstrap samples to calculate the standard errors of brain saliences for each LV. In this case, the assignment of the experimental conditions for all observations were maintained while each bootstrap was generated with replacement, i.e., the subject data were dropped one at a time and PLS was recalculated for each bootstrap sample. The confidence intervals are derived through bootstrap estimation, which is depicted in the LV brain-behaviour correlation profiles. The bootstrap ratio for each voxel is calculated as the voxel’s bootstrapped mean salience divided by its estimated standard error. Thus, higher BSR value represent more stable voxel saliences related to a given LV. In the present study, a bootstrap ratio (BSR) threshold of 3.00 (for a significance of p < 0.01) with a minimum spatial cluster size of 10 voxels was considered significant. We computed temporal brain scores to determine which time lags the task differences in a given LV were strongest (i.e., lags 2-5). Thus, it was for these peak voxel coordinates, which demonstrated maximal task differences, that were converted from MNI to Talairach space using the icbm2tal transform (Lancaster et al., 2007) as used in GingerAle 2.3 (Eickhoff et al., 2009). The cluster report for each LV was generated, which lists the brain regions involved in that particular LV, which meet the BSR and spatial cluster size threshold criteria. The cluster report also reveals additional ROI characteristics, including its BSR value, spatial extent, and the Brodmann area the coordinate falls under. Each ROI in the cluster report shows the peak coordinate of activation within the cluster. Peak coordinates from cerebellum and brainstem areas were removed from the cluster report because our fMRI acquisition did not completely acquire these regions.

Lastly, the B-PLS computes brain scores for each participant, which represents how strongly subject’s data reflect the LV observed. Thus, individuals with higher brain scores express the LV effect to a greater degree than subjects will lower brain scores. In order to verify our interpretations of the brain-behaviour patterns observed in each LV, we tested specific post-hoc LMER models of the brain scores. The fixed effects were the variables of Age, Sex, and Spatial Accuracy. In addition, to account for inter-individual variability within subjects tested across task conditions (i.e., eSE, eSH, rSE, rSH), we used Subject and Task Condition as random effects variables within the model. Significance was computed via the Satterthwaite approximation (at p ≤ 0.05) (using the R ‘lmerTest’ package, Kuznetsova et al., 2017).

## 3. Results

### 3.1. Behavioural Results

Table 1 shows mean and standard error values for demographic and fMRI behavioural measures, separated by Age Group and Sex. This includes group means for the CVLT, years of education, and retrieval accuracy scores (% correct) and reaction times (msec) for each event type. There were no significant differences in age or education level between sexes within each age group cohort. The LMER model testing the three-way interaction of Age, Sex, and Task Difficulty using spatial context retrieval accuracy as the dependent variable revealed a significant main effect of Age (beta = −0.0, SE = 0.01, t(123.73) =−2.21, p < 0.05), Task Difficulty (beta = – 0.04, SE =0.01, t(78) =−2.98, p < 0.05), and an interaction of Age*Task Difficulty (beta = −0.03, SE =0.01, t(78) = −2.01, p < 0.05). This interaction revealed that participants performed with significantly greater accuracy on the SE task compared to the SH task and that performance worsened on both tasks with increasing age (Figure 1).

**Figure 1.**
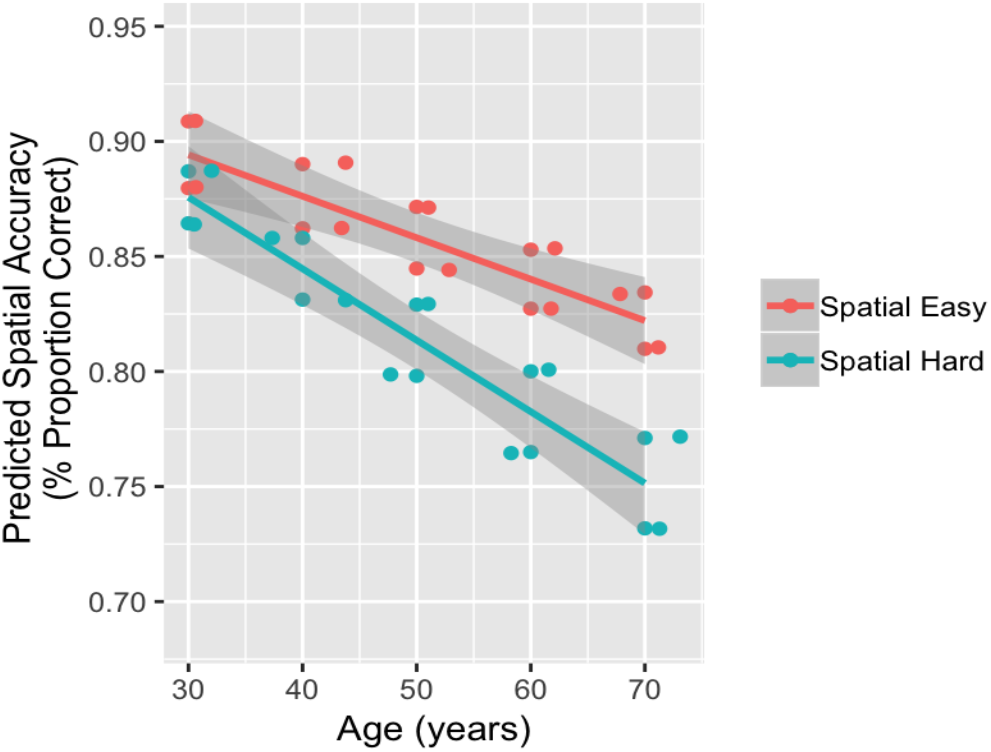
The partial effects plot demonstrating the two-way interaction of Age and Task Difficulty against Spatial Accuracy, showing worsening of memory performance with Age. Accuracy was higher for the SE task compared to the SH task. The shaded error bands represent the 95% confidence interval limits.

The LMER model testing the three-way interaction of Age, Sex, and Task Difficulty on Accuracy RT indicated main effects of Age (beta = 177.219, SE = 75.69, t(88.58) = 2.34, p < 0.05) and Task Difficulty (beta =137.15, SE = 37.82, t(78.0) = 3.63, p < 0.05). The main effect of Age demonstrated longer RT with greater age, whereas the main effect of Task Difficulty revealed that participants took longer to respond to the SH task compared to the SE task.

### 3.2. fMRI Results

The B-PLS analysis identified four significant latent variables (LVs; p < 0.05). Briefly, LV 1 (16.3% crossblock covariance), LV 3 (11.4%), and LV 4 (8.9%) identified sex differences in how age impacted memory-related brain activity at encoding, and/or retrieval, and in how brain activity related to performance. In contrast, LV 2 (14.2%) identified brain regions in which activity increased or decreased with age in both men and women, across encoding and retrieval (Age main effect; Supplementary Table 1 & Supplementary Figure 1), an effect that may be confounded by factors other than age-related changes in cognitive function (e.g., changes in CBF, CBV, vascular reactivity, and/or metabolism with age). Thus, this LV will not be further discussed because of the difficulties in disentangling all these factors that can impact the neurovascular coupling and subsequently affect the BOLD signal (D’Esposito, Deouell, & Gazzaley, 2003; C. L. Grady & Garrett, 2014; Handwerker, Gazzaley, Inglis, & D’Esposito, 2007; Kannurpatti, Motes, Rypma, & Biswal, 2010; Liu et al., 2013). Therefore, our results and discussion will focus specifically on sex differences in the effect of age and/or memory performance in memory-related brain activity. In the following sections, we present the results from LVs 1, 3, and 4 in detail.

#### 3.2.1. LV1

Figure 2A-B shows the singular image and correlation profiles for age and retrieval accuracy in men and women. Table 2 shows the local maxima for this LV. This LV only identified significant negative (blue coloured regions in the singular image) salience regions at the thresholds specified within bilateral inferior parietal cortex (IPC, peaking in left hemisphere), bilateral lateral prefrontal cortex (peaking in ventrolateral PFC [VLPFC]), right anterior-medial PFC and left parahippocampal gyrus (PHG). The correlation profile for age indicates that in men, age was not significantly correlated with activity in areas identified by this LV. In contrast, in women, advanced age was positively correlated with activity in negative salience regions at encoding and negatively correlated with activity in these regions during SH retrieval. Consistent with this interpretation, the post-hoc within sex LMER model of Age*Phase (4: eSE, eSH, rSE, rSH) interaction against brain scores revealed an Age*Phase interaction in women (p < 0.05) but not in men (p> 0.05). Therefore, this LV identified an Age*Phase (encoding/retrieval) interaction in women only.

**Figure 2.**
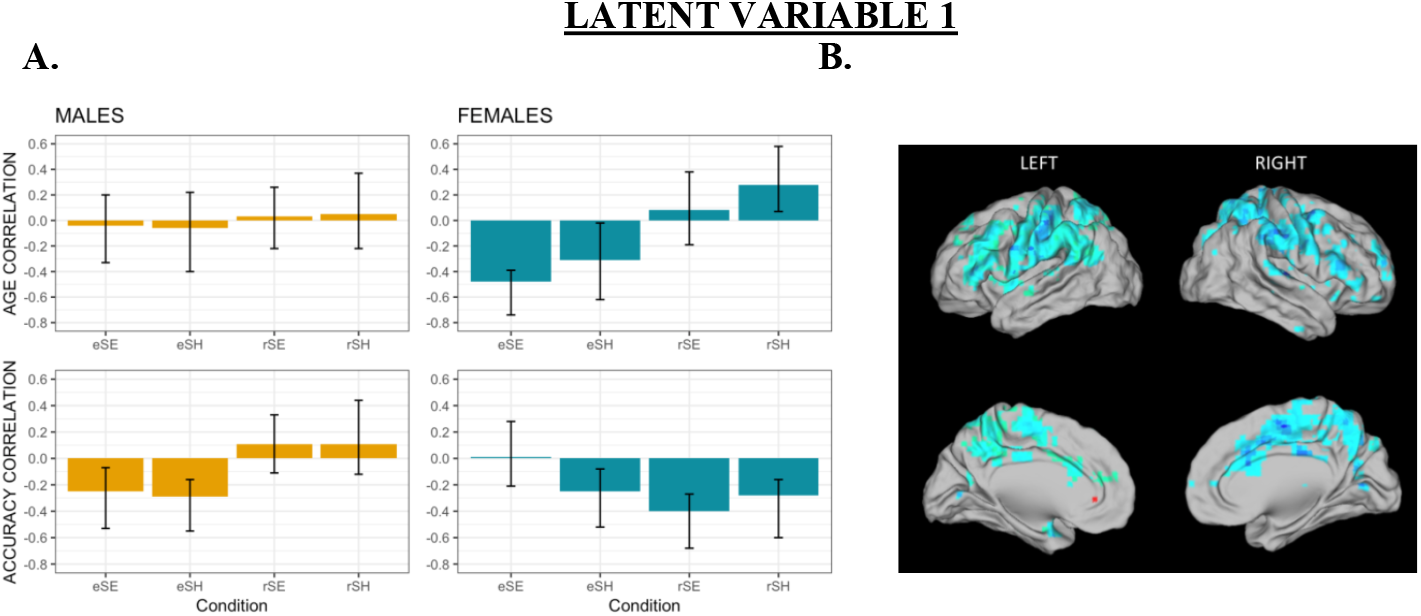
A) The brain-behaviour correlation profiles for LV1 for age and accuracy; B) The singular image for B-PLS LV1, threshold bootstrap ratio of ± 3.00, p < 0.001. Red brain regions reflect positive brain saliences and blue regions reflect negative brain saliences. Activations are presented on template images of the lateral and medial surfaces of the left and right hemispheres of the brain using Caret software (http://brainvis.wustl.edu/wiki/index.php/Caret:Download).

**Table 2.**
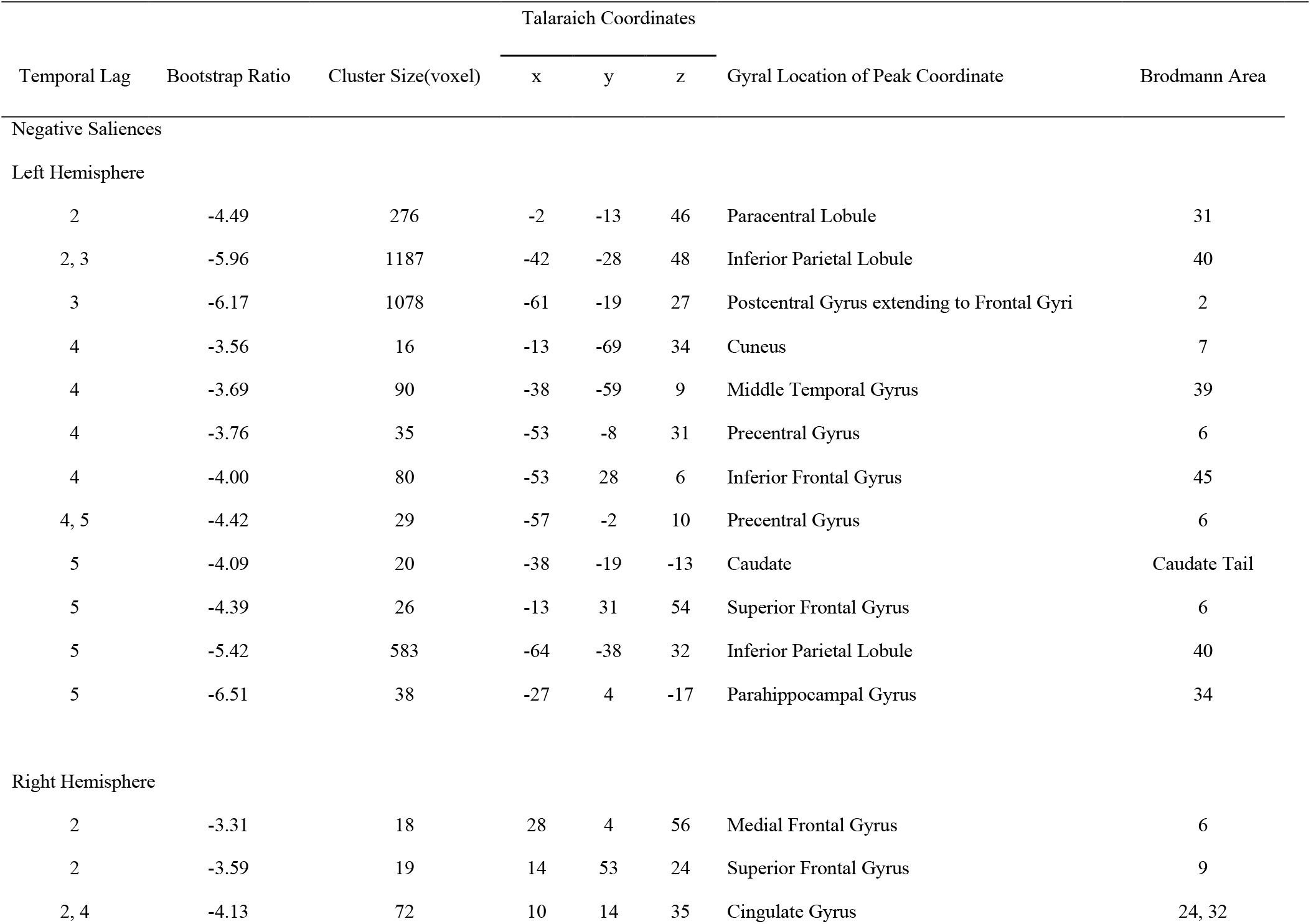
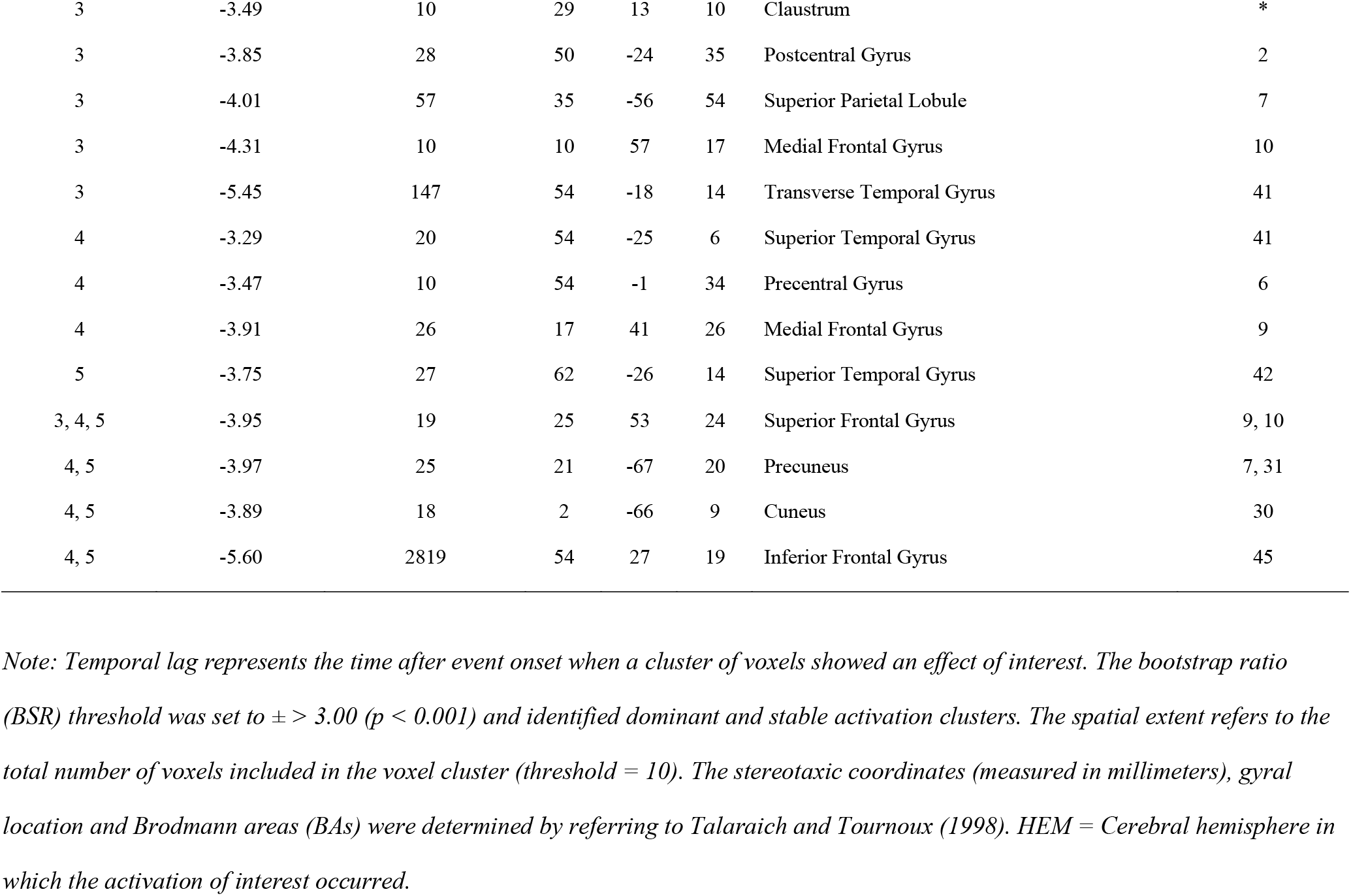
Local maxima for LV1

In relation to performance, the correlation profile revealed that men and women shared common encoding-related activity during the SH task, where increased encoding activity in negative salience regions was correlated with better subsequent retrieval accuracy. In addition, men showed that increased encoding activity in these same regions was related to better subsequent retrieval accuracy for the SH task. In women only, we observed that increased activity in negative salience regions during both retrieval tasks was positively correlated with retrieval accuracy. This is consistent with the post-hoc LMER model which revealed a significant Sex*Spatial Accuracy interaction (beta = −5.13, SE = 2.25, t(165.67) = −2.28, p < 0.05), indicating that both men and women showed decreasing brain scores with increasing spatial accuracy (Figure 3). This suggests a similar modulation of the LV pattern in both sexes, but that women contributed more heavily to this LV pattern compared to men.

**Figure 3.**
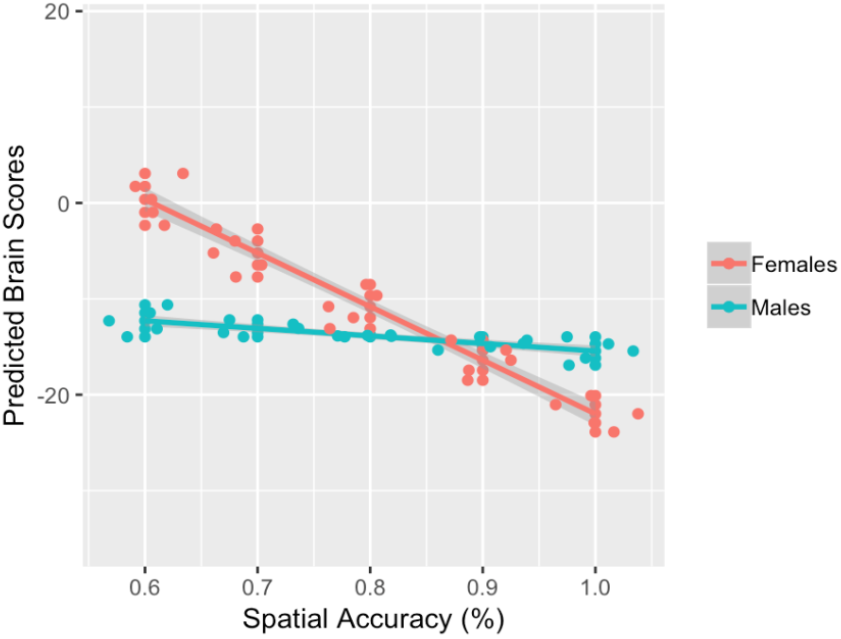
The partial effects plot demonstrating the interaction between Spatial Accuracy and Sex against brain scores (LV1). This effect shows that women contributed greater to the LV pattern observed compared to men except at a higher spatial accuracy. The shaded error bands represent the 95% confidence interval limits.

Therefore, in women age-related increases in left PHG, bilateral lateral PFC, right anterior PFC and parietal activity during SH encoding tasks was associated with subsequent memory. However, age-related decreases in activity within negative salience regions during SH retrieval in women was associated with worse retrieval accuracy. Compared to women, men showed no age-related changes directly linked to subsequent memory performance. Instead, men showed increased activity in these same fronto-parietal and PHG regions at encoding which was related to better subsequent memory for both tasks and was unrelated to age. These findings are supported by analyses of the brain scores demonstrating accuracy-related similarities between both sexes despite an Age*Phase interaction found in women but not men.

#### 3.2.2. LV3

Figure 4A-B shows the singular image and correlation profile for LV 3. Table 3 shows the local maxima of the regions involved in this LV. Unlike LV1, LV3 identified a distributed pattern of brain activity containing both negative *and* positive voxel saliences. This means that in relation to the brain-behaviour correlation profile, behavioural measures with negative correlations with task conditions are positively related to brain activity in negative voxel salience regions and negatively related to brain activity in positive voxel salience regions during the specified task conditions. Conversely, behavioural measures with positive correlations with task conditions are negatively related to negative voxel salience regions and positively related to positive voxel salience regions.

**Figure 4.**
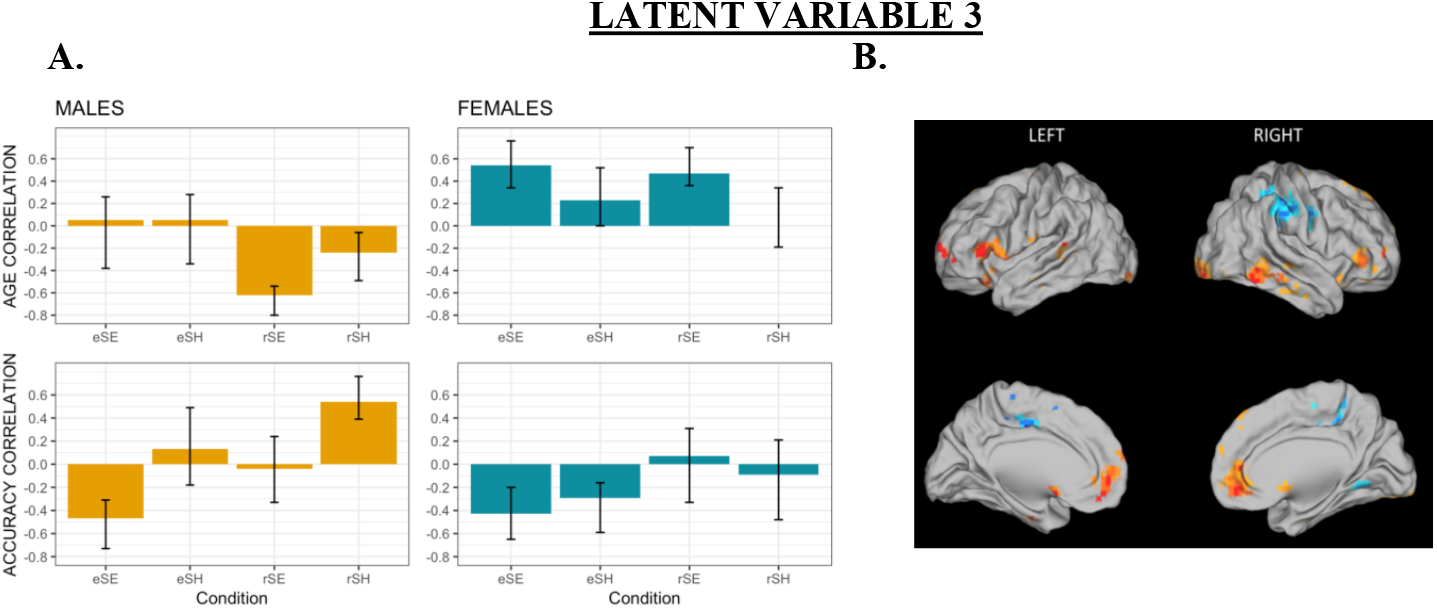
A) The brain-behaviour correlation profiles for LV3 for age and accuracy; B) The singular image for B-PLS LV3, threshold bootstrap ratio of ± 3.00, p < 0.001. Red brain regions reflect positive brain saliences and blue regions reflect negative brain saliences. Activations are presented on template images of the lateral and medial surfaces of the left and right hemispheres of the brain using Caret software (http://brainvis.wustl.edu/wiki/index.php/Caret:Download).

**Table 3.** Local maxima for LV3

| Temporal Lag | Bootstrap Ratio | Cluster Size (voxels) | Talarach Coordinates |  |  | Gyral Location | Brodmann Area |
| --- | --- | --- | --- | --- | --- | --- | --- |
|  |  |  | x | y | z |  |  |
| Positive Saliences |  |  |  |  |  |  |  |
| Left Hemisphere |  |  |  |  |  |  |  |
| 2 | 3.78 | 20 | -60 | -57 | -6 | Inferior Temporal Gyrus | 37 |
| 2, 5 | 4.08 | 21 | -13 | 27 | 61 | Superior Frontal Gyrus | 6 |
| 3 | 4.85 | 17 | -21 | -46 | 72 | Postcentral Gyrus | 7 |
| 3 | 3.76 | 21 | -31 | 47 | 8 | Middle Frontal Gyrus | 10 |
| 4 | 3.75 | 15 | -53 | -32 | 8 | Middle Temporal Gyrus | 22 |
| 5 | 5.16 | 49 | -1 | 3 | -6 | Anterior Cingulate | 25 |
| 4, 5 | 4.58 | 70 | -53 | 22 | -2 | Inferior Frontal Gyrus | 47 |
| 5 | 4.27 | 45 | -34 | 34 | -11 | Middle/Inferior Frontal Gyrus | 11/47 |
| 5 | 4.04 | 19 | -35 | -15 | 67 | Precentral Gyrus | 6 |
| 5 | 3.90 | 71 | -8 | 44 | -3 | Anterior Cingulate | 32 |
| 5 | 3.86 | 30 | -31 | -94 | -12 | Inferior Occipital Gyrus | 18 |
| 5 | 3.67 | 40 | -42 | -18 | -20 | Inferior Temporal Gyrus | 20 |
| 5 | 3.35 | 10 | -20 | 1 | 11 | Lentiform Nucleus | Putamen |
| Right Hemisphere |  |  |  |  |  |  |  |
| 3 | 4.70 | 57 | 58 | -46 | -10 | Inferior Temporal Gyrus | 20 |
| 3 | 4.65 | 31 | 39 | 0 | 63 | Middle Frontal Gyrus | 6 |
| 3 | 4.27 | 17 | 33 | 33 | -6 | Inferior Frontal Gyrus | 47 |
| 3 | 4.08 | 11 | 20 | -50 | 72 | Postcentral Gyrus | 7 |
| 2, 3, 5 | 4.80 | 134 | 17 | 15 | 64 | Superior Frontal Gyrus | 6, 8 |
| 2, 3, 4 | 4.72 | 189 | 7 | 47 | 5 | Anterior Cingulate | 32 |
| 4 | 4.58 | 33 | 51 | 35 | 8 | Inferior Frontal Gyrus | 46 |
| 4 | 3.85 | 10 | 44 | -12 | -7 | Insula | 13 |
| 4, 5 | 5.92 | 93 | 29 | -90 | -15 | Inferior Occipital/Fusiform Gyrus | 18 |
| 5 | 4.87 | 51 | 25 | 41 | -9 | Middle Frontal Gyrus | 11 |
| 5 | 4.40 | 23 | 47 | -20 | 0 | Superior Temporal Gyrus | 22 |
| 5 | 4.30 | 59 | 31 | -19 | 68 | Precentral Gyrus | 6 |
| 5 | 4.15 | 13 | 44 | 43 | 2 | Middle/Inferior Frontal Gyrus | 46 |
| 5 | 3.99 | 13 | 51 | -22 | -19 | Inferior Temporal Gyrus | 20 |
| Negative Saliences |  |  |  |  |  |  |  |
| Left Hemisphere |  |  |  |  |  |  |  |
| 3 | -4.40 | 61 | -16 | -20 | 42 | Cingulate Gyrus | 31 |
| 5 | -3.50 | 35 | -20 | -44 | 50 | Paracentral Lobule | 5 |
| 5 | -3.77 | 27 | -13 | -22 | 56 | Medial Frontal Gyrus | 6 |
| Right Hemisphere |  |  |  |  |  |  |  |
| 2, 3 | -4.59 | 330 | 39 | -4 | 26 | Precentral Gyrus | 6 |
| 4 | -3.89 | 53 | 39 | -24 | 39 | Postcentral Gyrus | 2 |
| 5 | -3.58 | 17 | 13 | -22 | 53 | Medial Frontal Gyrus | 6 |
| 5 | -4.29 | 301 | 17 | -39 | 40 | Cingulate Gyrus | 31 |

*Note: Temporal lag represents the time after event onset when a cluster of voxels showed an effect of interest. The bootstrap ratio (BSR) threshold was set to $\pm > 3.00$ ( $p < 0.001$ ) and identified dominant and stable activation clusters. The spatial extent refers to the total number of voxels included in the voxel cluster (threshold = 10). The stereotaxic coordinates (measured in millimeters), gyral location and Brodmann areas (BAs) were determined by referring to Talarach and Tournoux (1998). HEM = Cerebral hemisphere in which the activation of interest occurred.*

More specifically, in men, advanced age was positively correlated with activity in negative salience regions, and negatively correlated with activity in positive salience brain regions during retrieval. Specifically, older, compared to younger, men, exhibited greater activity in bilateral posterior cingulate (BA 31), and right postcentral gyrus extending into the temporoparietal junction – negative salience regions – at retrieval. In contrast, older, compared to younger, men exhibited *less* activity in bilateral lateral occipital, middle and superior-temporal cortices, bilateral VLPFC, and anterior cingulate (positive salience regions) at retrieval. In women, the inverse pattern of age-related activity was observed across encoding and retrieval phases of SE tasks. Thus, in women, advanced aged was related to greater activity in positive salience regions and decreased activity in negative salience regions during SE encoding and retrieval. However, when directly comparing age-related activity between both sexes as a function of task condition, men and women specifically showed the opposite pattern of brain activity during retrieval (SE task). The post-hoc LMER analysis on brain scores revealed a main effect of Age (beta = −2.91, SE = 1.09, t(91.42) = −2.67, p < 0.05) and an Age*Sex interaction (beta = 6.03, SE = 1.52, t(87.87) = 3.96, p < 0.05) which is consistent with the interaction observed at the LV level. Figure 5 depicts this interaction, which reveals that females have an age-related increase, whereas males have an age-related decrease, in brain scores. This suggests that the behavioural pattern of Age is more modulated by women compared to men in this LV.

**Figure 5.**
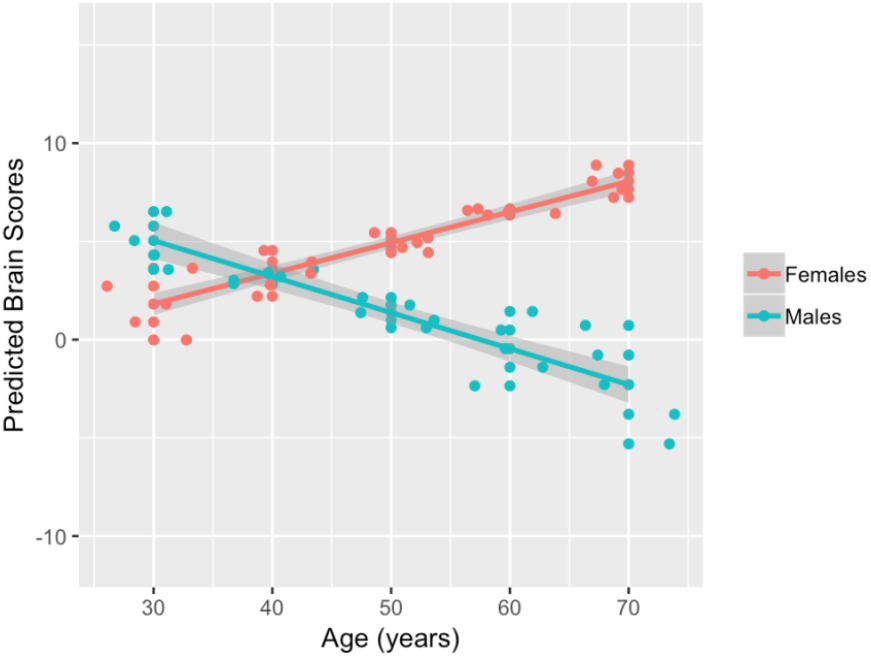
The partial effects plot demonstrating the interaction between Age and Sex against brain scores (LV3). This effect shows a greater age-related contribution of the LV pattern in women compared to men. The shaded error bands represent the 95% confidence interval limits.

In relation to performance, both sexes showed similar brain activity of regions at encoding of the SE task that was related to better subsequent memory. That is, in women, increased activity in negative salience regions, and decreased activity in positive salience regions, during encoding, was related to better subsequent memory. The same pattern of performance-related correlations was observed in men at encoding, but only for the SE task; however, the inverse pattern of performance-related correlations was observed in men at retrieval, for the SH task. Unlike men, women did not show significant performance related correlations at retrieval. The post-hoc LMER analysis did not reveal an effect or an interaction with Spatial Accuracy against brain scores. However, the post-hoc within sex LMER models testing Age*Spatial Accuracy against brain scores revealed a significant interaction in both sexes (p<0.05), which is consistent with the observed LV effect.

In summary, age-related differences in brain activity, at encoding in women and at retrieval in men, were negatively associated with subsequent memory performance. For example, in men, increased retrieval activity in positive salience regions and decreased retrieval activity in negative salience regions during SH tasks was related to better retrieval accuracy. However, with advanced age, men exhibited the opposite pattern of age-related differences in brain activity, which was related to poorer retrieval performance. In women, increased encoding-related activity in negative salience regions, and decreased encoding-related activity in positive salience-regions was related to better subsequent memory effects. Yet, older, compared to younger women, exhibited the opposite pattern of activity during SE encoding and retrieval tasks. Therefore, we observed sex differences as a function of task phase (i.e., encoding vs. retrieval) in how age correlated with memory-related brain activity. These differential patterns of age-related brain activity were related to worse memory performance in both sexes. Thus, older men and women’s poorer memory performance may be related to distinct differences in brain function with advance age.

#### 3.2.3. LV4

Figure 6A-B shows the singular image and correlation profile for LV 4. Table 4 shows the local maxima of the regions involved in this LV. The correlation profile indicates that in men, increasing age was positively correlated with increased encoding-related activity in negative salience regions, including activity in anterior cingulate and right DLPFC. In contrast, in men, increasing age was *negatively* correlated with encoding-related activity in positive salience regions, including: bilateral lateral occipital and temporal cortices, and bilateral parahippocampal gyrus (PHG) extending to right hippocampus. In women, a main effect of age was observed across encoding and retrieval that was in the opposite direction of that observed in men. Specifically, in women, increasing age was positively correlated with activity in positive salience regions and negatively correlated with activity in negative salience regions during encoding and retrieval. The post-hoc LMER analysis which directly tested this Age*Sex interaction against brain scores found this interaction to be significant (beta = 3.89, SE = 1.30, t(73.94) =2.99, p < 0.05). Figure 7 depicts this interaction, which revealed that at a younger age, males have higher brain scores compared to females, but the inverse was true in older age.

**Figure 6.**
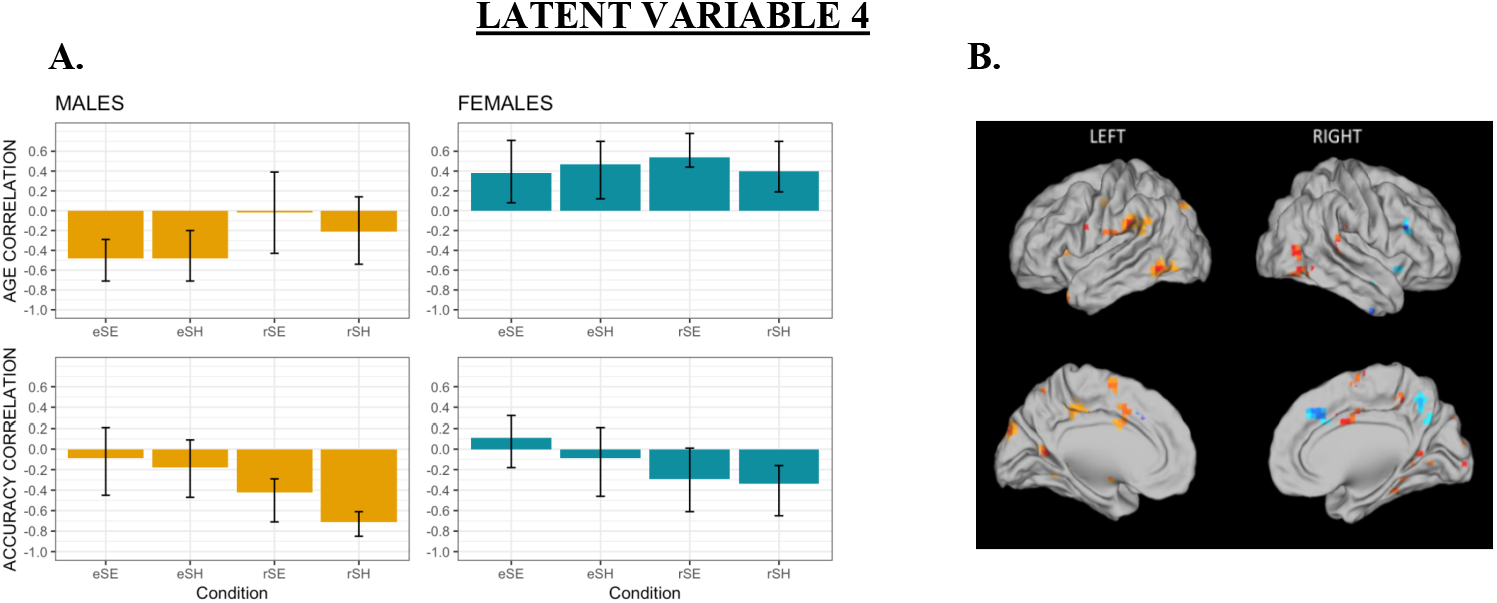
A) The brain-behaviour correlation profiles for LV4 for age and accuracy; B) The singular image for B-PLS LV4, threshold bootstrap ratio of ± 3.00, p < 0.001. Red brain regions reflect positive brain saliences and blue regions reflect negative brain saliences. Activations are presented on template images of the lateral and medial surfaces of the left and right hemispheres of the brain using Caret software (http://brainvis.wustl.edu/wiki/index.php/Caret:Download).

**Figure 7.**
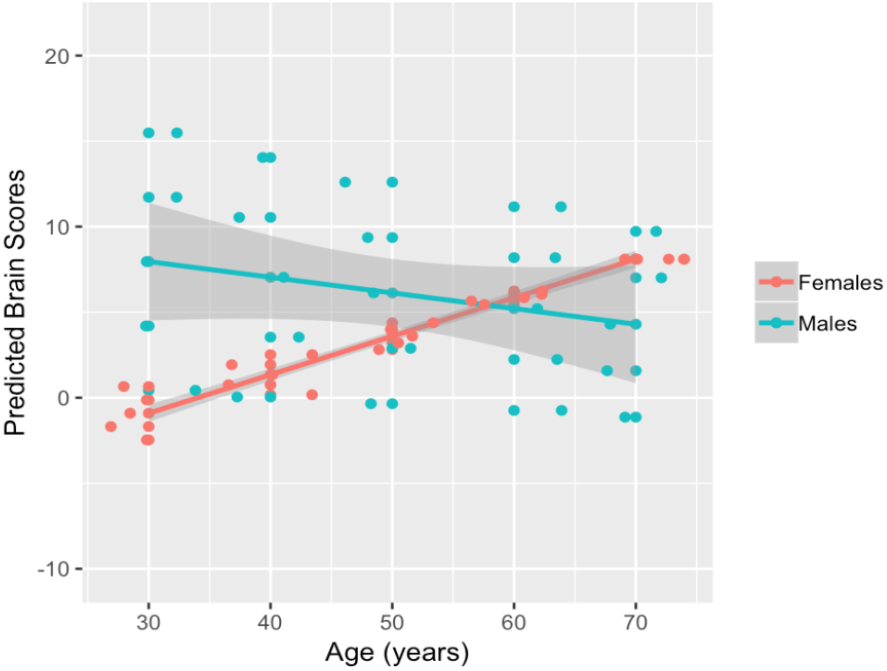
The partial effects plot demonstrating the significant interaction of Age and Sex against brain scores, indicating a greater age-related modulation of brain scores in women compared to men (LV4). The shaded error bands represent the 95% confidence interval limits.

**Table 4.**
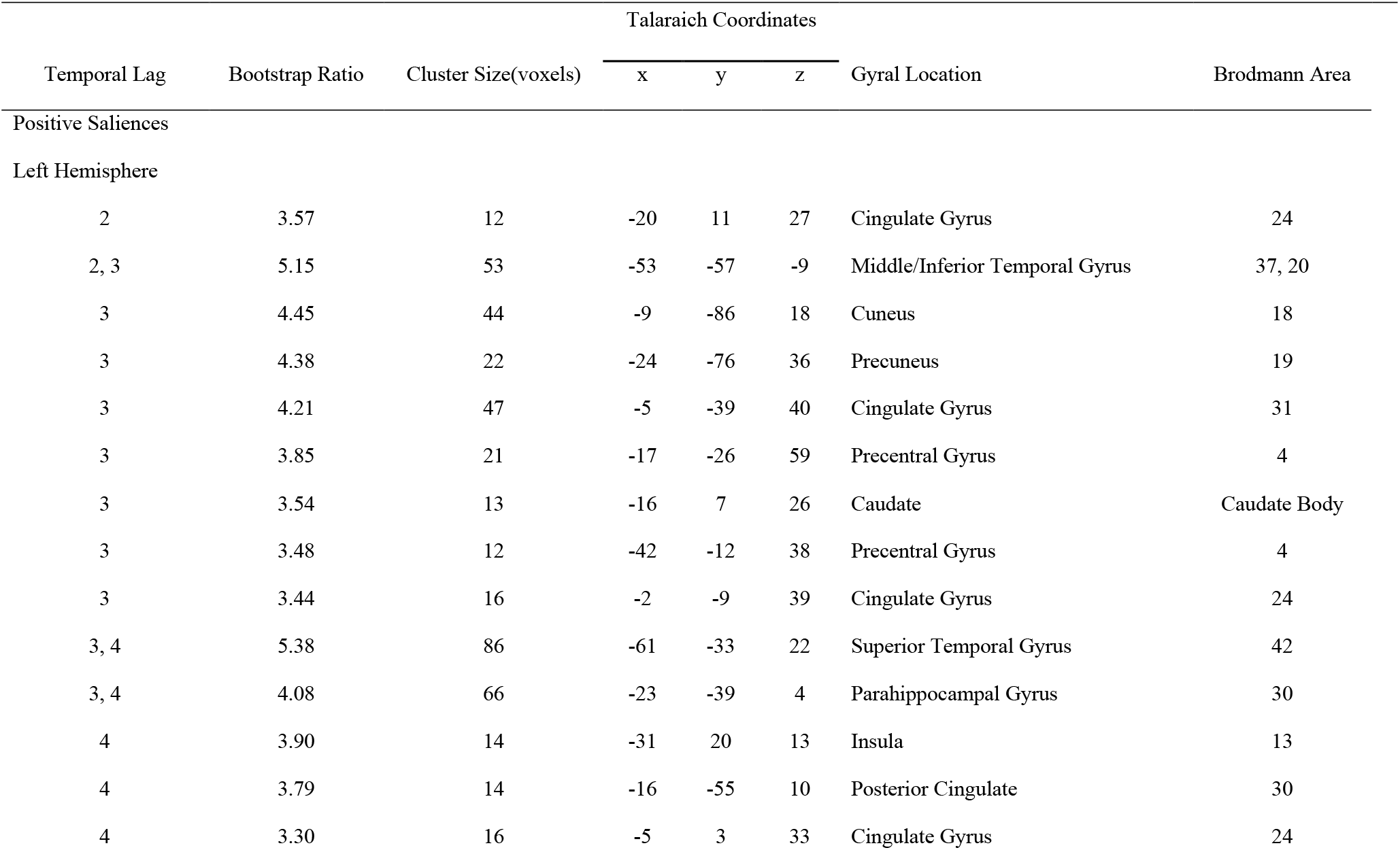
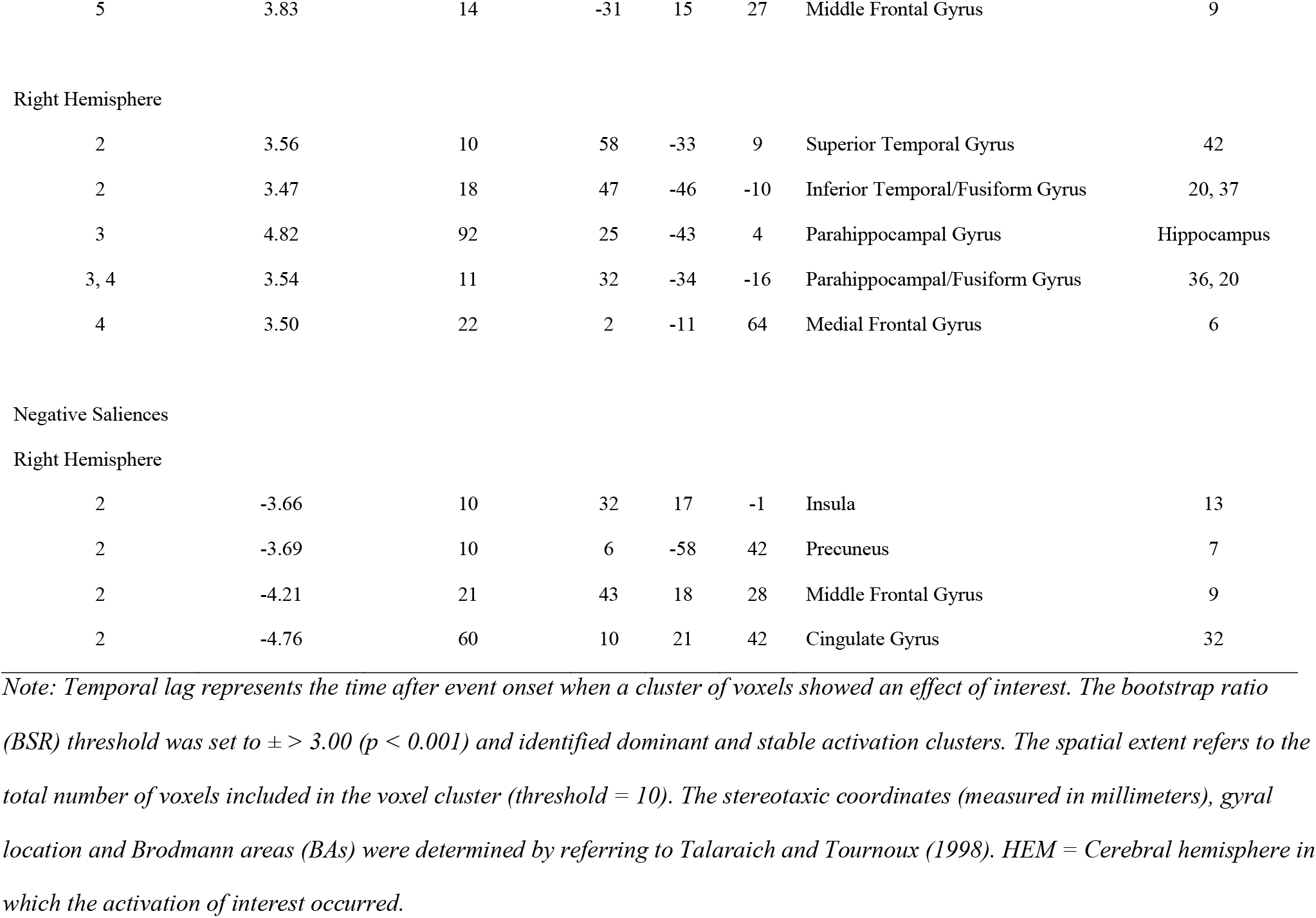
Local maxima for LV4

| Temporal Lag | Bootstrap Ratio | Cluster Size(voxels) | Talaraich Coordinates |  |  | Gyral Location | Brodmann Area |
| --- | --- | --- | --- | --- | --- | --- | --- |
|  |  |  | x | y | z |  |  |
| Positive Saliences |  |  |  |  |  |  |  |
| Left Hemisphere |  |  |  |  |  |  |  |
| 2 | 3.57 | 12 | -20 | 11 | 27 | Cingulate Gyrus | 24 |
| 2, 3 | 5.15 | 53 | -53 | -57 | -9 | Middle/Inferior Temporal Gyrus | 37, 20 |
| 3 | 4.45 | 44 | -9 | -86 | 18 | Cuneus | 18 |
| 3 | 4.38 | 22 | -24 | -76 | 36 | Precuneus | 19 |
| 3 | 4.21 | 47 | -5 | -39 | 40 | Cingulate Gyrus | 31 |
| 3 | 3.85 | 21 | -17 | -26 | 59 | Precentral Gyrus | 4 |
| 3 | 3.54 | 13 | -16 | 7 | 26 | Caudate | Caudate Body |
| 3 | 3.48 | 12 | -42 | -12 | 38 | Precentral Gyrus | 4 |
| 3 | 3.44 | 16 | -2 | -9 | 39 | Cingulate Gyrus | 24 |
| 3, 4 | 5.38 | 86 | -61 | -33 | 22 | Superior Temporal Gyrus | 42 |
| 3, 4 | 4.08 | 66 | -23 | -39 | 4 | Parahippocampal Gyrus | 30 |
| 4 | 3.90 | 14 | -31 | 20 | 13 | Insula | 13 |
| 4 | 3.79 | 14 | -16 | -55 | 10 | Posterior Cingulate | 30 |
| 4 | 3.30 | 16 | -5 | 3 | 33 | Cingulate Gyrus | 24 |
| 5 | 3.83 | 14 | -31 | 15 | 27 | Middle Frontal Gyrus | 9 |
| Right Hemisphere |  |  |  |  |  |  |  |
| 2 | 3.56 | 10 | 58 | -33 | 9 | Superior Temporal Gyrus | 42 |
| 2 | 3.47 | 18 | 47 | -46 | -10 | Inferior Temporal/Fusiform Gyrus | 20, 37 |
| 3 | 4.82 | 92 | 25 | -43 | 4 | Parahippocampal Gyrus | Hippocampus |
| 3, 4 | 3.54 | 11 | 32 | -34 | -16 | Parahippocampal/Fusiform Gyrus | 36, 20 |
| 4 | 3.50 | 22 | 2 | -11 | 64 | Medial Frontal Gyrus | 6 |
| Negative Saliences |  |  |  |  |  |  |  |
| Right Hemisphere |  |  |  |  |  |  |  |
| 2 | -3.66 | 10 | 32 | 17 | -1 | Insula | 13 |
| 2 | -3.69 | 10 | 6 | -58 | 42 | Precuneus | 7 |
| 2 | -4.21 | 21 | 43 | 18 | 28 | Middle Frontal Gyrus | 9 |
| 2 | -4.76 | 60 | 10 | 21 | 42 | Cingulate Gyrus | 32 |

*Note: Temporal lag represents the time after event onset when a cluster of voxels showed an effect of interest. The bootstrap ratio (BSR) threshold was set to $\pm > 3.00$ ( $p < 0.001$ ) and identified dominant and stable activation clusters. The spatial extent refers to the total number of voxels included in the voxel cluster (threshold = 10). The stereotaxic coordinates (measured in millimeters), gyral location and Brodmann areas (BAs) were determined by referring to Talarach and Tournoux (1998). HEM = Cerebral hemisphere in which the activation of interest occurred.*

In relation to retrieval accuracy, the correlation profile indicates that in both men and women, increased retrieval-related activity in negative salience regions (right DLPFC and anterior cingulate), and decreased retrieval-related activity in positive salience regions (left DLPFC, bilateral PHG, and lateral occipito-temporal cortices) was related to better retrieval accuracy. Consistent with this main effect of Spatial Accuracy observed in this LV, we tested for and found a main effect of Spatial Accuracy against brain scores (beta = −1.70, SE = 0.60, t(241.79) = −2.84, p < 0.05), which indicated that increasing spatial accuracy was associated with lower brain scores.

In summary, in women, age-related increases in brain activity in positive salience regions during retrieval was related to lower retrieval accuracy, particularly during SH retrieval tasks. Since men did not exhibit age-related differences in activity at retrieval, the brain-performance accuracy correlations observed at retrieval, were unrelated to age in men.

## 4. Discussion

In the current fMRI study, we investigated how age and biological sex were associated with brain activity during the encoding and retrieval of spatial context memory. We examined whether age-related differences in brain activity during a memory task were similar among women and men. To this aim, we used between group multivariate B-PLS to test for sex differences in the effect of age, memory performance and Age*Memory performance on encoding- and retrieval-related brain activity. The behavioural analysis of retrieval accuracy and RT measures obtained during fMRI tasks indicated there was a main effect of age: increasing age was related to declines in retrieval accuracy and increases in RT on spatial context memory tasks. This result is consistent with the prior results observed in our larger adult lifespan sample (Ankudowich et al., 2016, 2017) and with previous adult lifespan studies of context/source memory (e.g., Cansino, 2009; M. K. Johnson, Hashtroudi, & Lindsay, 1993). In addition, participants took longer and performance accuracy declined on the hard vs. easy version of the task, demonstrating an effect of task load on performance with aging, again, replicating previous findings (Ankudowich et al., 2016; Kwon et al., 2016; Spaniol & Grady, 2012). We did not observe a significant main effect of biological sex, nor did we observe a significant Age*Sex interaction in either retrieval accuracy or RT.

Our behavioural results are in contrast to prior studies that have shown women outperform men on a variety of episodic memory tasks: such as, verbal stimuli (Herlitz et al., 1997; Kimura & Harshman, 1984; Ragland et al., 2000), negative emotional stimuli (Young et al., 2013), face stimuli (Keightley et al., 2006; Sommer et al., 2013; Yonker et al., 2003) and verbal paired associated memory (Bender et al., 2010). The absence of an effect of sex, nor an age-by-sex interaction, in memory performance may be explained by the nature of the episodic memory task we used. This is the first study to our knowledge to explore sex differences in face-location spatial context memory. Past studies have shown that on average, men outperform women on spatial tasks, and women perform better than men on face and emotional memory tasks (Herlitz et al., 1997; Lejbak, Crossley, & Vrbancic, 2011; Sommer et al., 2013; Weiss et al., 2003). Thus, the null effect of sex on spatial context memory performance for face-location associations may have emerged because the stimuli used were non-verbal and required both spatial and facial/emotional stimuli processing. Alternatively, it is possible that when female and male participants are matched on education and neuropsychological test measures, as they were in the current study, sex differences in episodic memory are no longer apparent.

Although we did not observe a significant effect of Sex, or an Age*Sex interaction on spatial context retrieval accuracy and RT, our fMRI results identified sex differences in age-related patterns of brain activity during successful spatial context encoding and retrieval, and sex differences in performance-related patterns of brain activity during retrieval. Overall, our findings suggest that the neural correlates of age-related spatial context memory decline differ in women compared to men. Below we discuss our findings in greater detail.

### 4.1. Performance-related patterns of brain activity: Similarities and differences between the sexes

Our findings suggest that generally women and men engaged similar brain regions at encoding and retrieval to support memory performance. This observation is consistent with the behavioural results indicating there were no significant sex differences in memory performance. In both sexes, activity in right anterior-medial, bilateral dorsal and lateral PFC and left PHG during spatial context encoding positively correlated with better subsequent memory (LV1). Similarly, activity in right dorsolateral PFC (DLPFC) during spatial context retrieval positively correlated with retrieval accuracy (LV4). These results are consistent with prior literature showing that successful episodic encoding is associated with medial temporal and lateral PFC activity, and successful episodic retrieval is associated with right DLPFC activity (Blumenfeld & Ranganath, 2007; Hayama & Rugg, 2009; Murray & Ranganath, 2007; Simons & Spiers, 2003; Staresina, 2006).

LV1 also indicated that increased inferior parietal activity, extending into PFC, at retrieval was positively related to memory performance in women. Previously, we examined age and performance-related differences in brain activity using a larger lifespan cohort where sample size for the two sexes were not matched (Ankudowich et al., 2017). We did not consider biological sex in this earlier analysis. Results from this analysis indicated that inferior parietal activity during both encoding and retrieval reflected performance, rather than age, effects. Interestingly, by considering sex in the current analysis we observe that frontal and parietal activation at encoding supported subsequent performance in both sexes, but frontal and parietal activation at retrieval was only positively correlated to memory performance in women. Prior studies have highlighted the importance of frontoparietal regions in mediating top-down cognitive control of memory-related medial temporal functions at encoding and retrieval (Cabeza et al., 2003; Cabeza, Dolcos, Graham, & Nyberg, 2002; Dulas & Duarte, 2014; Grady, 2008; Mitchell & Johnson, 2009). The current study suggests that the controlled relational encoding of face-location associations benefitted subsequent memory in both sexes, but the recapitulation of these processes at retrieval only supported memory performance in women.

We also found that activity in bilateral lateral temporal cortices, posterior lateral occipital cortices, anterior cingulate and ventrolateral PFC (VLPFC) during *encoding* (LV3), was negatively correlated with subsequent memory in both sexes. However, in men, increased activity in anterior lateral occipital, middle and superior temporal cortices, and VLPFC at retrieval supported retrieval accuracy on hard spatial context memory tasks. In addition, we also found that activity in more posterior lateral temporal and occipital regions during retrieval (LV4) was negatively related to memory performance in both sexes. These findings are generally consistent with our prior analysis of this study, in which sex differences were not investigated (Ankudowich et al., 2017). In both the previous and current analyses, we found that activity in lateral temporal and lateral posterior occipital areas during encoding and retrieval was negatively correlated with performance, and positively correlated with age. In the current study, we found that this positive correlation with age was primarily observed in women (see below).

Activity in lateral occipital-temporal cortices and VLPFC at encoding has been associated with less-specific, semantic processing of visual stimuli, including faces (Demb et al., 1995; Kirchhoff, Anderson, Barch, & Jacoby, 2012). This suggests that semantic processing during the encoding of face-location associations results in poorer memory in the current study but controlled relational encoding of face-location associations (discussed above) supported subsequent memory. This observation is consistent with recent findings showing that orienting subjects to semantic associations was detrimental to subsequent memory and suggests that semantically associating stimuli at encoding may interfere with successful episodic encoding (Long & Kahana, 2017). Interestingly, our LV3 retrieval effects suggest that in men engaging VLPFC, lateral occipital (anterior) and temporal cortices supported retrieval accuracy.

In general, our findings suggest that sex differences in performance-related activity were present at retrieval compared to encoding. In women, encoding *and* retrieval success was related to the engagement of lateral frontoparietal-related cognitive control processes, and to relational mnemonic processes associated with the PHG (LV1 and LV4). In men these same processes were important for successful spatial context encoding success, but successful retrieval was also related to the retrieval of semantic associations.

Our findings demonstrate that both sexes engaged different sets of brain regions and related cognitive processes to support memory at retrieval. To our knowledge, very few studies have specifically explored functional sex differences at episodic retrieval. One such study by Nyberg et al (2000) found functional sex differences at memory retrieval in a younger adult sample. Similar to our findings, Nyberg et al found increased activity in bilateral inferior temporal cortex in males compared to females at retrieval. However, their sex difference findings in bilateral inferior parietal cortex are contrary to what we observed in our results. Specifically, Nyberg et al found that women had reduced inferior parietal cortical activity compared to men. This contradicts our findings of sex differences at retrieval, where men and women showed the opposite pattern of results in fusiform and IPC activity. These differences may be because these studies used different experimental tasks. That is, Nyberg et al. combined data from three different episodic memory tasks, which differed in terms of the type of stimuli encoded (i.e., words, sentences, landscapes), the way in which participants had to encode the stimuli (i.e., silent reading, intentional encoding), and the modality in which the stimuli were presented at encoding (i.e., visual vs. auditory). Moreover, their retrieval task was a yes/no recognition task of the stimuli, whereas our retrieval task was more explicit in that participants had to select the face that they saw either to the left/right at encoding (depending on the retrieval cue). Taken together, these studies showed that participants recruited brain regions typically involved at retrieval, and suggest that the specific experimental tasks used might help explain differences in retrieval-related brain activity across study findings. Below we discuss sex differences in the effect of age on memory related brain activity.

### 4.2. Sex differences in the effect of age on encoding and retrieval related activity

We observed sex differences in the effect of age on frontal-parietal (LV1), medial temporal activity (LV1; LV4); lateral occipital-temporal (LV3; LV4) and VLPFC (LV3) activity during encoding and retrieval. Specifically, with advanced age women exhibited increased activity in lateral occipital-temporal cortices across encoding and retrieval (LV3; LV4); particularly for easy spatial context memory tasks. Women also exhibited age-related increases in right PHG during encoding and retrieval (LV4), and age-related increases in frontal-parietal and left PHG during encoding (LV1). Thus, with advanced age women exhibited a general increase in brain activity in a variety of regions at encoding and retrieval.

In contrast, women exhibited age-related decreases in right DLPFC activity across encoding and retrieval (LV4); and in bilateral frontal-parietal and left PHG during hard spatial context retrieval (LV1). Age-related decreases in these regions may reflect an age-related deficit in function, particularly at retrieval, given that activation of these regions at retrieval was related to better memory performance (discussed above) (Rajah & D’Esposito, 2005). This suggests that in women, age-related reductions in spatial context memory may be related to differences in PFC, parietal and medial temporal activity at retrieval.

Also, in women, activity in lateral occipital-temporal at encoding and retrieval, in VLPFC at encoding was negatively correlated with memory performance. Thus, the observation that with advanced age, women increased activity in these brain regions suggests that older women may have engaged in sub-optimal strategies during encoding and retrieval (Mitchell & Johnson, 2009). In contrast, given that the engagement of frontal-parietal and left PHG activity at encoding and retrieval was related to better memory performance in women; the age-related increase in these regions at encoding may reflect functional compensation (Cabeza et al., 2018).

In men, advanced age was related to decreased activity in lateral occipital-temporal and right PHG at encoding (LV4), and in lateral occipital-temporal and VLPFC activity at retrieval (LV3). Given that activation of these brain regions during hard spatial context retrieval tasks was related to better memory performance in men, these sex specific decreases in lateral occipital-temporal and VLPFC activity suggests there may be functional deficits in these regions in older men. In contrast, men exhibited age-related increases in precentral and posterior cingulate activity during retrieval and in right DLPFC and cingulate during encoding. These patterns of age-related increase were not beneficial to performance in men. Therefore, in men we did not observe age-related differences in lateral frontal-parietal and left PHG activity (LV1) and activation of these regions at encoding supported subsequent memory across all ages. In both men and women, activation of right DLPFC during retrieval supported memory and activation of lateral occipital-temporal and VLPFC at retrieval support memory only in men. However, the pattern of age-related activation in these regions at encoding and retrieval was not beneficial to memory performance.

## 5. Conclusions

Our results point to several significant sex differences in how age impacts memory-related brain function. First, that age-related increases in lateral frontal-parietal and left PHG activity at encoding were specific to women and directly impacted memory success. Second, there were pronounced sex differences in the impact of age on occipital, temporal and VLPFC activity, however in both sexes the pattern of age-related difference in these regional activations were similarly detrimental to task performance. In other words, although women exhibited a generalized age-related increase in brain activity in these areas across encoding and retrieval, and men exhibited age-related decreases in activity within these regions only at retrieval; in both sexes, these age-related differences negatively impacted memory performance. Third, age-related deficits in spatial context memory are primarily related to altered brain activity at retrieval in both sexes, but the nature of that age-related differences in activation was not the same in women and men. This implies that the neural correlates of spatial context memory decline with age differs in men and women, and the brain regions that support successful memory function in late life also differ in women and men. Moreover, we were able to verify the patterns observed at the LV level through post-hoc LMER analyses using brain scores generated for each participant across task conditions. In conclusion, our study showed that men and women showed both functional similarities and differences, which were modulated by age, in context memory performance. Understanding these similarities and differences are critical in providing important insight as to why there are sex differences in memory-related disorders and how treatment interventions should be tailored in the realm of aging and dementia.

### 5.1. Caveats

The present investigation explored sex similarities and differences in episodic/context memory across the lifespan. There are limitations of the study design and methodology that prevent us from having a more comprehensive understanding of biological sex effects on memory and aging. First, it is important to consider how hormonal differences between men and women and age-related hormonal changes within sex might influence our interpretation of the results. For example, past studies have shown the phase of the menstrual cycle (linking changes in estradiol and progesterone) have significantly contributed to differences in spatial memory performance in women (Doreen Kimura & Hampson, 1994). In addition, midlife in women is associated with declines in 17β -estradiol, a known neurocognitive hormone that might contribute to performance differences in midlife compared to other age groups (e.g., Jacobs et al., 2016b; Morrison, Brinton, Schmidt, & Gore, 2006; Rentz et al., 2017). To minimize our variability, we removed MA women with HRT in the present study design (see Methods). However, an understanding of hormonal changes in aging is important given that women report subjective memory deficits particularly during menopause transition (Weber & Mapstone, 2009). Finally, it is important to consider hormonal differences, such as age-related declines in testosterone (a hormone linked to spatial memory), which may selectively impact the cognitive aging process in both sexes (Fabbri et al., 2016; Harman, Metter, Tobin, Pearson, & Blackman, 2001). Importantly, the influence of societal gender roles in addition to stress differences (e.g., Pruessner, 2018) will meaningfully contribute to our understanding of sex differences in memory and aging. Thus, a more holistic approach in understanding sex differences in memory and aging, which takes into consideration biological and sociocultural differences, might shed more light on why women are at greater risk for memory-related disorders, such as AD.

## Acknowledgments

This work was supported by Canadian Institute of Health Sciences (CIHR) Project Grant #M0P126105 and Alzheimer’s Society of Canada Research Grant # 1435 awarded to M.N. Rajah, and by the Natural Science and Engineering Research Council (NSERC) Graham Bell Canada Graduate Scholarship-Doctoral (CGS D) awarded to S. Subramaniapillai. We thank Dr. A. Almey for her editorial help with the Discussion.

---

**Supplementary Figure 1.**
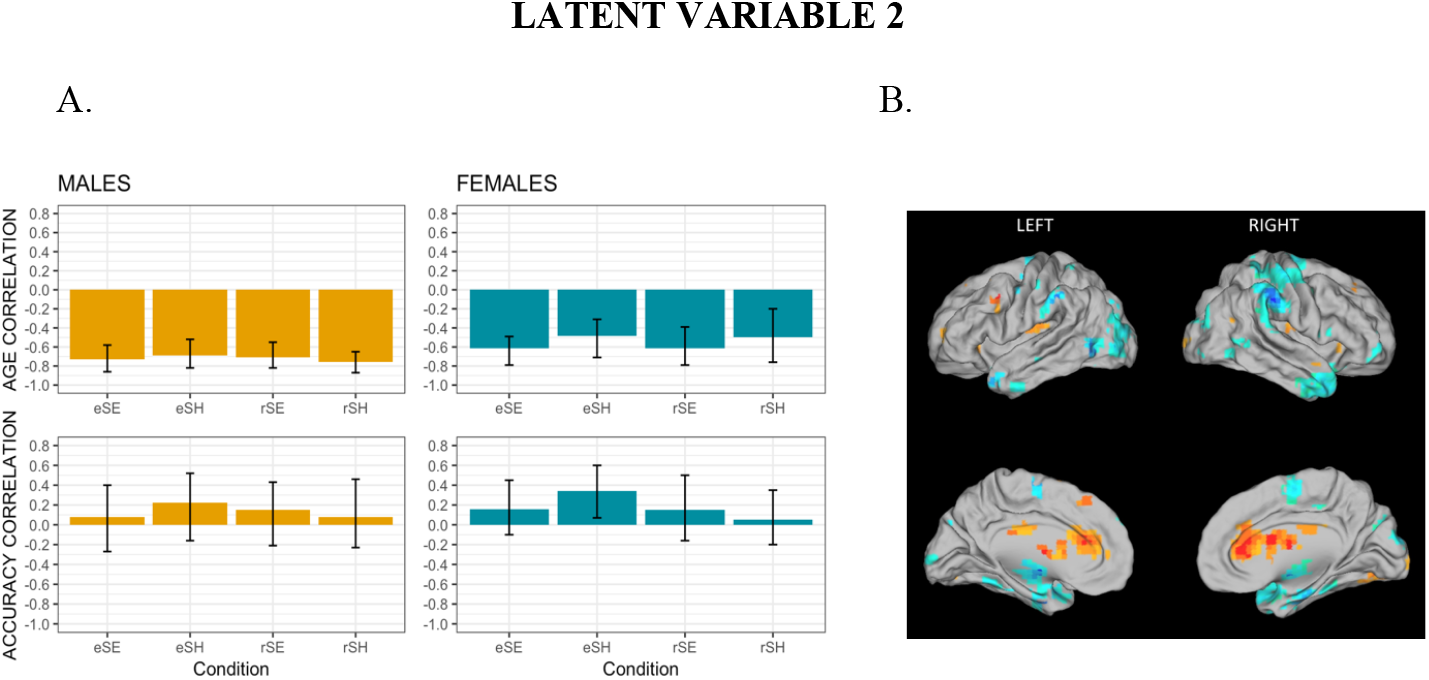
A) The brain-behaviour correlation profiles for LV2 for age and accuracy; B) The singular image for B-PLS LV2, threshold bootstrap ratio of ± 3.00, p < 0.001. Red brain regions reflect positive brain saliences and blue regions reflect negative brain saliences. Activations are presented on template images of the lateral and medial surfaces of the left and right hemispheres of the brain using Caret software (http://brainvis.wustl.edu/wiki/index.php/Caret:Download).

**Supplementary Table 1.**
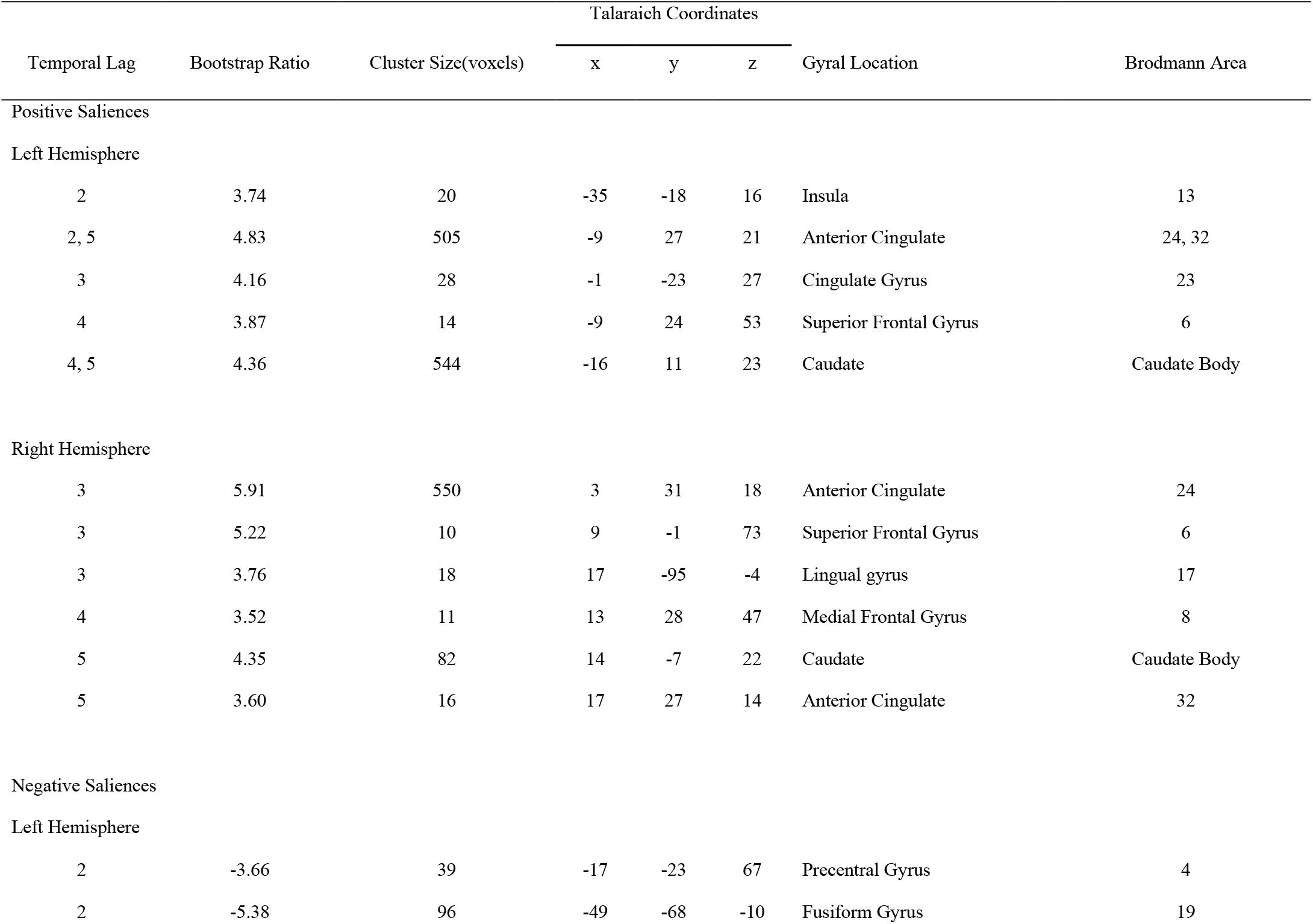
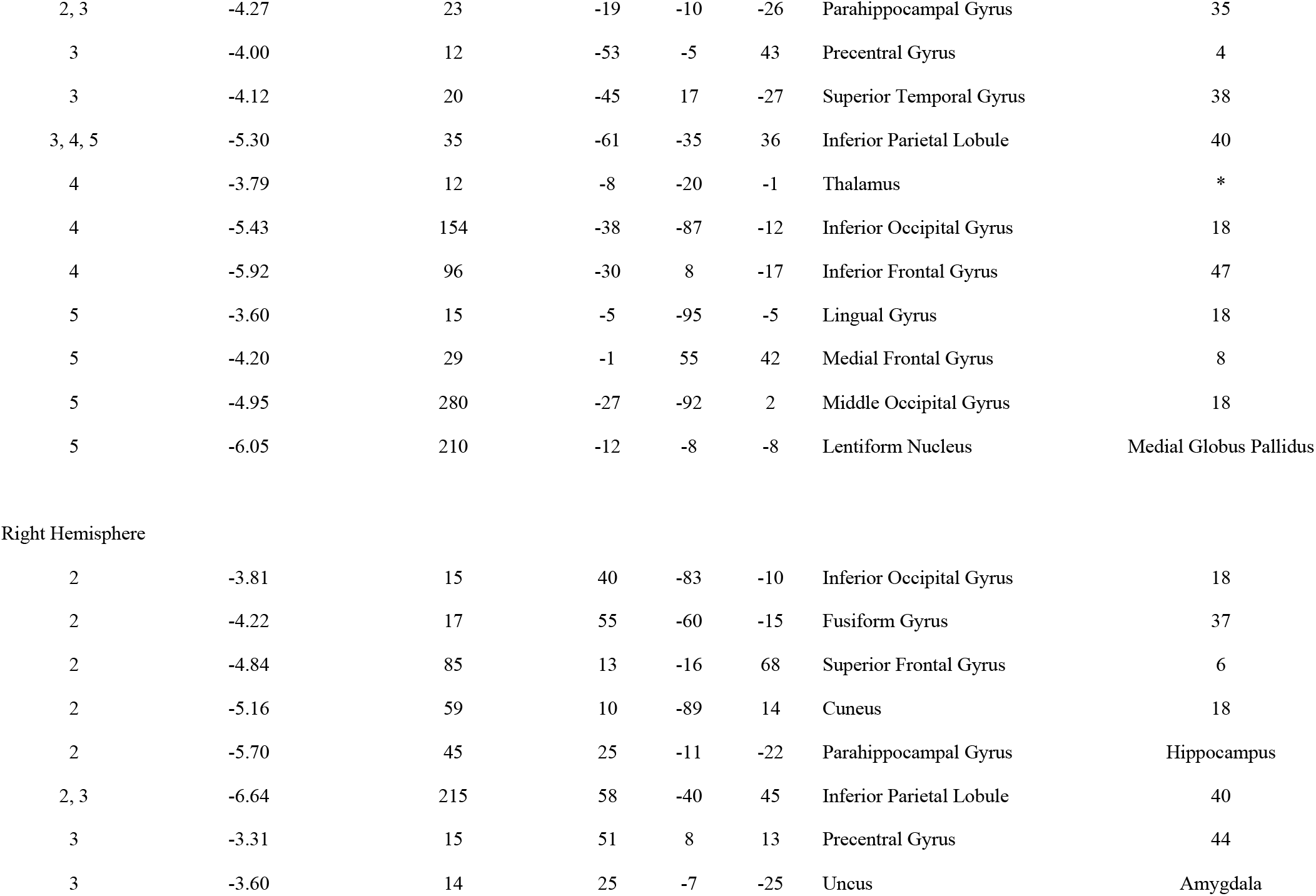
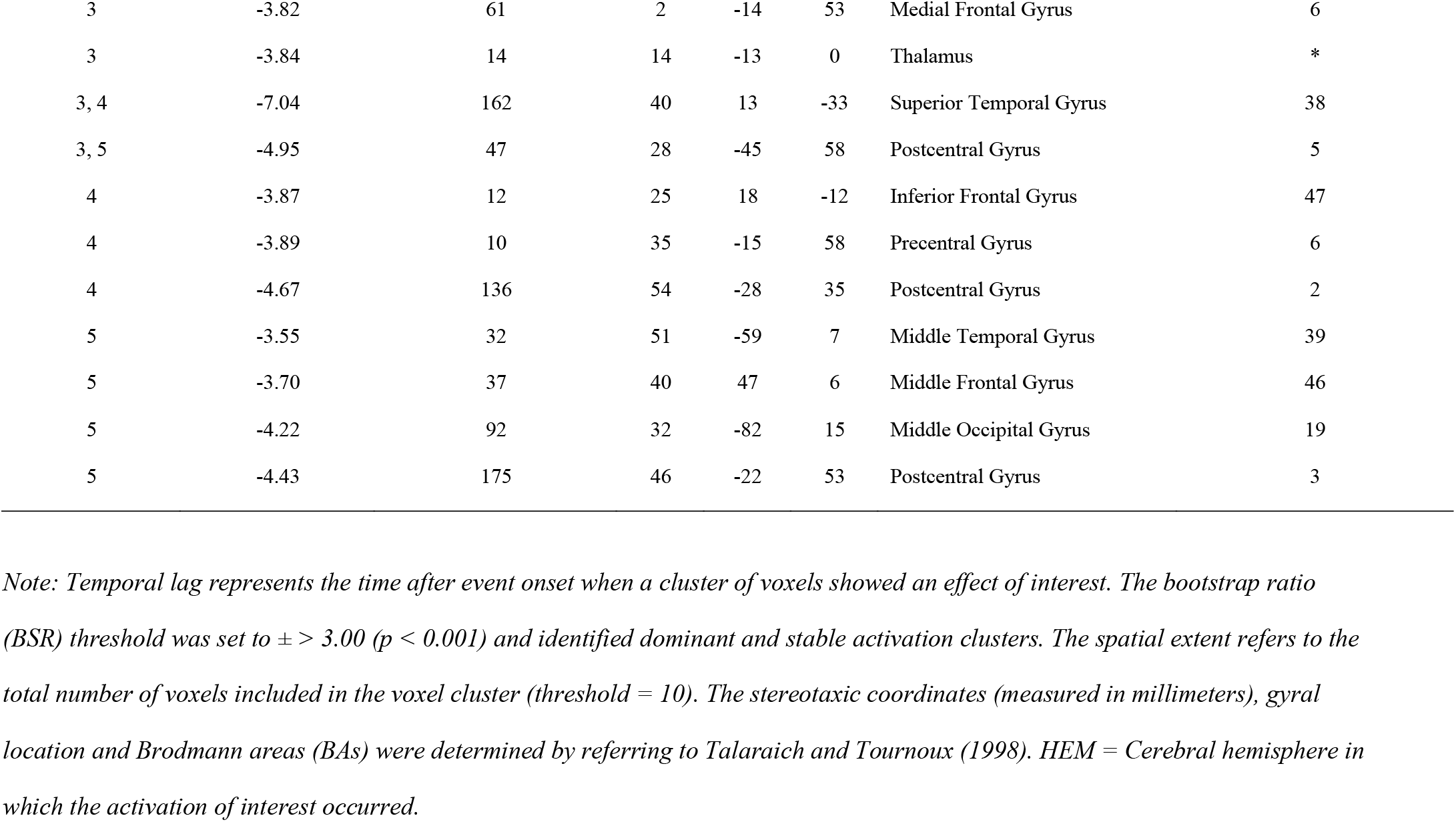
Local maxima for LV2

| Temporal Lag | Bootstrap Ratio | Cluster Size(voxels) | Talaraich Coordinates |  |  | Gyral Location | Brodmann Area |
| --- | --- | --- | --- | --- | --- | --- | --- |
|  |  |  | x | y | z |  |  |
| Positive Saliences |  |  |  |  |  |  |  |
| Left Hemisphere |  |  |  |  |  |  |  |
| 2 | 3.74 | 20 | -35 | -18 | 16 | Insula | 13 |
| 2, 5 | 4.83 | 505 | -9 | 27 | 21 | Anterior Cingulate | 24, 32 |
| 3 | 4.16 | 28 | -1 | -23 | 27 | Cingulate Gyrus | 23 |
| 4 | 3.87 | 14 | -9 | 24 | 53 | Superior Frontal Gyrus | 6 |
| 4, 5 | 4.36 | 544 | -16 | 11 | 23 | Caudate | Caudate Body |
| Right Hemisphere |  |  |  |  |  |  |  |
| 3 | 5.91 | 550 | 3 | 31 | 18 | Anterior Cingulate | 24 |
| 3 | 5.22 | 10 | 9 | -1 | 73 | Superior Frontal Gyrus | 6 |
| 3 | 3.76 | 18 | 17 | -95 | -4 | Lingual gyrus | 17 |
| 4 | 3.52 | 11 | 13 | 28 | 47 | Medial Frontal Gyrus | 8 |
| 5 | 4.35 | 82 | 14 | -7 | 22 | Caudate | Caudate Body |
| 5 | 3.60 | 16 | 17 | 27 | 14 | Anterior Cingulate | 32 |
| Negative Saliences |  |  |  |  |  |  |  |
| Left Hemisphere |  |  |  |  |  |  |  |
| 2 | -3.66 | 39 | -17 | -23 | 67 | Precentral Gyrus | 4 |
| 2 | -5.38 | 96 | -49 | -68 | -10 | Fusiform Gyrus | 19 |
| 2, 3 | -4.27 | 23 | -19 | -10 | -26 | Parahippocampal Gyrus | 35 |
| 3 | -4.00 | 12 | -53 | -5 | 43 | Precentral Gyrus | 4 |
| 3 | -4.12 | 20 | -45 | 17 | -27 | Superior Temporal Gyrus | 38 |
| 3, 4, 5 | -5.30 | 35 | -61 | -35 | 36 | Inferior Parietal Lobule | 40 |
| 4 | -3.79 | 12 | -8 | -20 | -1 | Thalamus | * |
| 4 | -5.43 | 154 | -38 | -87 | -12 | Inferior Occipital Gyrus | 18 |
| 4 | -5.92 | 96 | -30 | 8 | -17 | Inferior Frontal Gyrus | 47 |
| 5 | -3.60 | 15 | -5 | -95 | -5 | Lingual Gyrus | 18 |
| 5 | -4.20 | 29 | -1 | 55 | 42 | Medial Frontal Gyrus | 8 |
| 5 | -4.95 | 280 | -27 | -92 | 2 | Middle Occipital Gyrus | 18 |
| 5 | -6.05 | 210 | -12 | -8 | -8 | Lentiform Nucleus | Medial Globus Pallidus |
| Right Hemisphere |  |  |  |  |  |  |  |
| 2 | -3.81 | 15 | 40 | -83 | -10 | Inferior Occipital Gyrus | 18 |
| 2 | -4.22 | 17 | 55 | -60 | -15 | Fusiform Gyrus | 37 |
| 2 | -4.84 | 85 | 13 | -16 | 68 | Superior Frontal Gyrus | 6 |
| 2 | -5.16 | 59 | 10 | -89 | 14 | Cuneus | 18 |
| 2 | -5.70 | 45 | 25 | -11 | -22 | Parahippocampal Gyrus | Hippocampus |
| 2, 3 | -6.64 | 215 | 58 | -40 | 45 | Inferior Parietal Lobule | 40 |
| 3 | -3.31 | 15 | 51 | 8 | 13 | Precentral Gyrus | 44 |
| 3 | -3.60 | 14 | 25 | -7 | -25 | Uncus | Amygdala |
| 3 | -3.82 | 61 | 2 | -14 | 53 | Medial Frontal Gyrus | 6 |
| 3 | -3.84 | 14 | 14 | -13 | 0 | Thalamus | * |
| 3, 4 | -7.04 | 162 | 40 | 13 | -33 | Superior Temporal Gyrus | 38 |
| 3, 5 | -4.95 | 47 | 28 | -45 | 58 | Postcentral Gyrus | 5 |
| 4 | -3.87 | 12 | 25 | 18 | -12 | Inferior Frontal Gyrus | 47 |
| 4 | -3.89 | 10 | 35 | -15 | 58 | Precentral Gyrus | 6 |
| 4 | -4.67 | 136 | 54 | -28 | 35 | Postcentral Gyrus | 2 |
| 5 | -3.55 | 32 | 51 | -59 | 7 | Middle Temporal Gyrus | 39 |
| 5 | -3.70 | 37 | 40 | 47 | 6 | Middle Frontal Gyrus | 46 |
| 5 | -4.22 | 92 | 32 | -82 | 15 | Middle Occipital Gyrus | 19 |
| 5 | -4.43 | 175 | 46 | -22 | 53 | Postcentral Gyrus | 3 |

*Note: Temporal lag represents the time after event onset when a cluster of voxels showed an effect of interest. The bootstrap ratio (BSR) threshold was set to $\pm > 3.00$ ( $p < 0.001$ ) and identified dominant and stable activation clusters. The spatial extent refers to the total number of voxels included in the voxel cluster (threshold = 10). The stereotaxic coordinates (measured in millimeters), gyral location and Brodmann areas (BAs) were determined by referring to Talarach and Tournoux (1998). HEM = Cerebral hemisphere in which the activation of interest occurred.*

